# Mechanism of Gating and Isoform-Specific Inhibition in Renal CLC Chloride Channels

**DOI:** 10.64898/2026.02.17.706469

**Authors:** Chih-Ta Chien, Briana L. Sobecks-Doherty, Alexander S. Powers, Anindita Das, Jürgen Kreiter, Chloe N. Barry, Muyuan Chen, Andrew Hinman, Camille F. Petrakian, Natasa Trifkovic, Brianna Williams, Chase A.P. Wood, Mengyuan Xu, Ron O. Dror, Wah Chiu, Merritt Maduke

## Abstract

Hyponatremia is a prevalent disorder marked by excess water retention and substantial morbidity, motivating interest in the CLC-Ka chloride channel as a therapeutic target. Selectively inhibiting CLC-Ka without affecting the closely related CLC-Kb is essential for preventing serious side effects. However, developing isoform-selective inhibitors has been challenging because most small molecules do not distinguish between CLC-Ka and CLC-Kb, and the basis for selectivity in the few known exceptions remains unclear. The small molecule BIM1 preferentially inhibits CLC-Ka over CLC-Kb, providing an opportunity to dissect isoform-specific pharmacology. To investigate this mechanism, we determined cryo-EM structures of BIM1 and BIM15, a related nonselective analog, bound to a CLC-K variant engineered to match the human CLC-Ka binding pocket. Structural and computational analyses reveal that inhibition and isoform selectivity are anchored by interactions with a conserved lysine, with surrounding binding-site residues subtly tuning the local electrostatic environment to promote or disfavor these contacts. These analyses further identify a dynamic extracellular loop that intermittently occludes the shared pathway accessing the inhibitor-binding site and pore. Bound BIM15 engages this gating loop more extensively than BIM1, suggesting that differential loop engagement contributes to inhibitor selectivity, a prediction validated by mutagenesis. Because loop dynamics block the pore, we examined the structural impact of Ca²⁺, which favors opening, and found the gating loop ordered and withdrawn from the pathway. Together, these findings define how binding-site microenvironments and gating-loop dynamics shape isoform-specific inhibition and pore access in CLC-K channels.

**Significance Statement:** Hyponatremia is a major clinical problem with limited therapeutic options. The kidney chloride channel CLC-Ka is an attractive drug target, but its high sequence identity to CLC-Kb has hindered the development of isoform-selective inhibitors needed for safe therapy. A low-micromolar CLC-Ka–selective inhibitor had been identified, providing a foothold for drug development, but the structural basis of its selectivity was unknown. Here, by integrating cryo-EM structures with molecular dynamics simulations, we define the inhibitor-binding site and reveal the mechanism that enables preferential CLC-Ka inhibition. We further show that a dynamic extracellular loop functions as a gating element shaping inhibitor access and engagement. These findings establish a mechanistic foundation for developing improved treatments for hyponatremia.

## Introduction

CLC (“Chloride channel”) proteins constitute a functionally heterogeneous family of plasma-membrane ion channels and intracellular transporters. By precisely regulating ion flux, they orchestrate diverse physiological processes, ranging from electrical excitability to cellular homeostasis (1, 2). In the kidney, two channel isoforms, CLC-Ka and CLC-Kb, play key roles in transepithelial salt and water homeostasis (3–6). CLC-Ka, found in the thin ascending limb of the nephron, establishes and sustains the osmotic gradient that is necessary for water reabsorption and the production of concentrated urine. CLC-Kb, expressed in the thick ascending limb and distal convoluted tubule, provides a key pathway for chloride (Cl^−^) to leave kidney cells and return to the blood. Loss of CLC-Kb function causes Bartter syndrome type III, a severe salt wasting disorder (7), and CLC-Kb has been suggested as a target for antihypertensive therapies (3, 8). In contrast, CLC-Ka is a potential target for treating hyponatremia. Hyponatremia, defined as low serum sodium and osmolality, requires a treatment that increases water excretion. Because CLC-Ka mediates Cl^−^ efflux in the thin ascending limb, the site where the medullary osmotic gradient that drives collecting-duct water reabsorption is established (9), its inhibition is expected to blunt this gradient and thereby increase water excretion (10, 11). Consistent with this mechanism, targeted disruption of CLC-Ka dissipates the medullary concentration gradient (12) and produces water diuresis (13).

In addition to their renal roles, CLC-Ka and CLC-Kb are co-expressed in the inner ear, where their redundant functions are essential for hearing. A major challenge in developing drugs targeting CLC-Ka or CLC-Kb is the high sequence identity (91%) between the two homologs. Drug selectivity is paramount not only to target the distinct therapeutic indications of CLC-Ka and CLC-Kb, but also because simultaneous inhibition of both channels results in Bartter syndrome type IV – a condition characterized by extreme salt wasting and deafness (14–16). Here, we address the key problem of how small molecules achieve selective inhibition of CLC-Ka over CLC-Kb.

Chloride-channel pharmacology has a troubled history, with classic “chloride-channel inhibitors” lacking the precision needed to distinguish between closely related channel subtypes or even unrelated ion channels (17). An oligomeric stilbene disulfonate was found to potently inhibit CLC-Ka with >100-fold discrimination against CLC-Kb, showing that selective CLC-Ka inhibition is feasible (18); however, that molecule is not drug-like and not tractable for synthetic optimization. Using insights from studies on low-potency inhibitors (11, 19), Liantonio and colleagues leveraged structure-activity relationship (SAR) analysis to develop MT-189, a low-micromolar CLC-Ka inhibitor with ∼3-fold selectivity over CLC-Kb (8, 20). Building on this foundation, we designed BIM1, which retains low-micromolar potency against CLC-Ka while boosting selectivity over CLC-Kb to twenty-fold (21). Docking simulations of BIM1 into CLC-Ka homology models implicated N68 and K165 in potency, but the marked variability in the identities and orientations of BIM-interacting residues across docking solutions precluded a definitive binding-pocket map (21) (Fig. 1). An improved understanding of BIM1 binding interactions and the molecular basis of its homolog specificity is critical for designing next-generation analogs with enhanced potency and preserved CLC-Ka/CLC-Kb selectivity, suitable for evaluation in preclinical studies.

**Figure 1.**
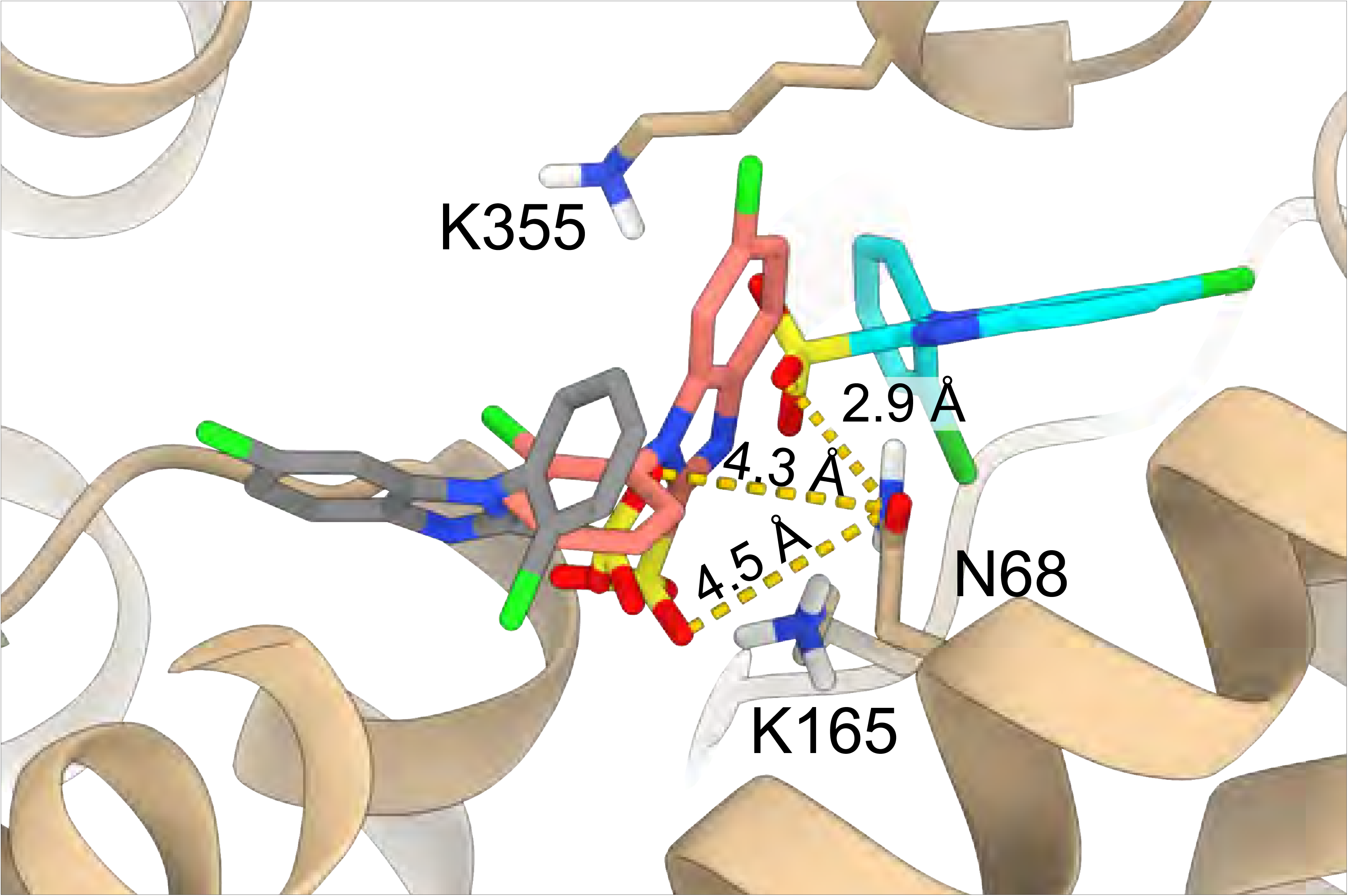
Variability in BIM1 orientation within the CLC-Ka binding pocket from prior computational docking studies. Three representative docking poses (21) are shown (in grey, salmon, and cyan) to illustrate the variability in BIM1 orientation within the pocket. Key residues N68, K165, and K355 are indicated, and distances from the closest oxygen of BIM1 to NZ atom of N68 – a major determinant of selectivity – are specified. In the present work, we introduced S68N and R355K into bovine CLC-K to reproduce the human CLC-Ka sequence at these positions, called “bCLC-Ka” this modified construct is referred to as bCLC-Ka throughout the manuscript.

## Results and Discussion

### Experimental system

We chose the bovine CLC-K homolog (bCLC-K) for structure-determination because its apo cryo-EM structure has been solved, providing established expression and purification protocols (22). Bovine CLC-K is a single paralog that shares 84% sequence identity with the two human CLC-K isoforms. In the computationally predicted BIM binding pocket (Fig. 1), two residues differ between bCLC-K and human CLC-Ka (hCLC-Ka) (Fig. S1), so we introduced S68N and R355K substitutions to make the bovine sequence match hCLC-Ka specifically at those positions; we refer to this engineered channel as bCLC-Ka. Bovine do not have separate CLC-Ka and CLC-Kb isoforms. Using two-electrode voltage clamp (TEVC) recordings of *Xenopus* oocytes, we found that bCLC-Ka exhibits BIM1 and BIM15 sensitivity comparable to hCLC-Ka (Fig. 2, Fig. S2). A difference between the two channels is that bCLC-Ka lacks the small gating relaxations displayed by hCLC-Ka at negative voltages. This gating behavior matches that of hCLC-Ka when expressed in HEK293 cells (23) and is intermediate between the mild hyperpolarization-activation of hCLC-Ka and the mild depolarization-activation of hCLC-Kb expressed in oocytes (21). Although the basis for the different voltage dependence of hCLC-Ka in oocytes versus HEK293 cells remains unknown, multiple inhibitors exhibit similar potency on hCLC-Ka channels in *Xenopus* oocyte and mammalian expression systems (23), supporting that these gating differences do not affect inhibitor efficacy. Thus, despite modest gating differences, the inhibitor sensitivity of bCLC-Ka is sufficiently similar to that of hCLC-Ka to justify its use for structure determination.

**Figure 2.**
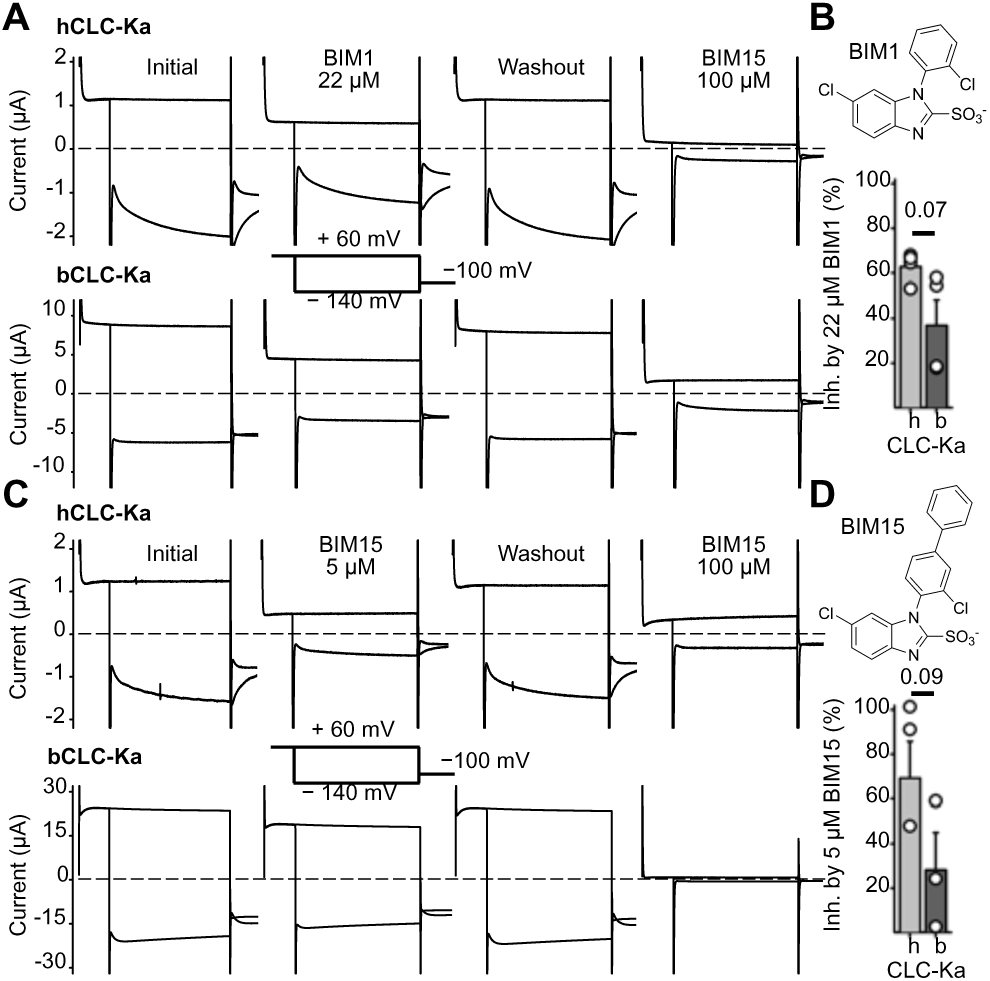
bCLC-Ka is sensitive to inhibition by BIM1 and BIM15 (*A*) TEVC recordings of oocytes overexpressing hCLC-Ka (upper traces) and bCLC-Ka (lower traces). Each set of traces show currents under four conditions: initial, after 22 µM BIM1, after BIM1 washout, and after subsequent application of 100 µM BIM15. The voltage protocol is depicted between the second set of traces. (*B*) Depiction of BIM1 chemical structure and summary data for inhibition of hCLC-Ka and bCLC-Ka by 22 µM BIM1. Bars represent the mean ± S.E.M; circles represent individual inhibition values (n = 4 biological replicates). Inhibition was calculated as described in Materials and Methods. The P-value was determined using an unpaired t-test. (*C*) TEVC recordings of oocytes overexpressing hCLC-Ka and bCLC-Ka, as in panel A except testing inhibition by 5 µM BIM15 in place of 22 µM BIM1. (*D*) Depiction of BIM15 chemical structure and summary data for inhibition of hCLC-Ka and bCLC-Ka by 5 µM BIM15. Bars represent the mean ± S.E.M; circles represent individual inhibition values (n = 3 biological replicates). Inhibition was calculated as described in Materials and Methods. The P-value was determined using an unpaired t-test.

### Sulfonate position in the BIM1-bound bCLC-Ka structure

We first determined the apo structure of bCLC-Ka in nanodiscs to establish the channel architecture in a native-like membrane environment. Using an adapted bCLC-K preparation protocol (22), we obtained a 3.6 Å reconstruction (Fig. S3) that closely matches the previously reported bCLC-K structure (PDB 5TQQ), which has been interpreted as adopting an open-like arrangement in comparison to CLC transporter structures (22), with an RMSD of 0.9 Å. We then determined the BIM1-bound structure at 2.8 Å resolution (Fig. S4). Its overall architecture matches the apo structure (RMSD = 0.6 Å), and BIM1 occupies the predicted binding site. The pore pathway is not significantly changed, aside from BIM1 physically blocking access from the extracellular side (Fig. 3*A*, Fig. S5*B,E*). A further distinction is the clear electron density consistent with Cl^−^ at the S_ext_ and S_cen_ sites (24), whereas no comparable density is present in the apo map (Fig. S6).

**Figure 3.**
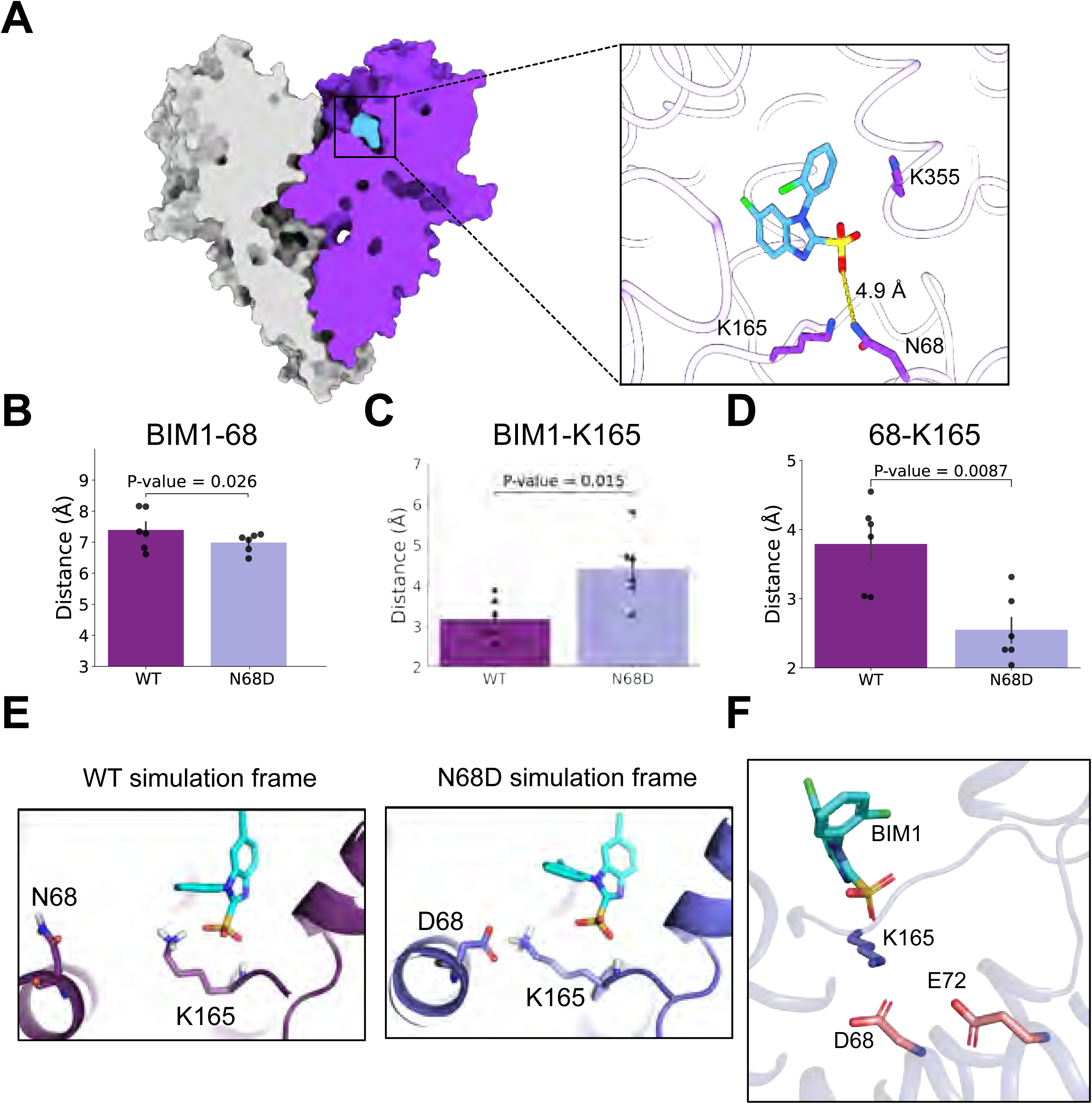
Mechanism of BIM1 CLC-K isoform selectivity. *(A)* Cryo-EM structure of bCLC-Ka bound to BIM1. The zoomed-in view of the binding pocket shows the distance between the side chain nitrogen of N68 and closest oxygen of the BIM sulfonate group. (*B*) N68 does not directly interact with the BIM sulfonate group. Bar graphs show the average distances between residue 68 (OD1 or OD2/ND2 atom) and BIM1 (closest sulfonate oxygen atom) across multiple simulations. (*C*) K165-BIM1 distances increase with N68D mutation. Distances are between the closest H atom on K165 and the closest sulfonate oxygen on BIM1. (*D*) K165-65 distances decrease with N68D mutation. Distances are between the closest H atom on K165 and the closest side chain oxygen/nitrogen of N68 or D68. For panels B-D, average distances for individual simulations are shown as black dots, with black bars indicating the SEM, and P-values were assessed using the non-parametric Mann–Whitney U test. (*E*) Representative MD frames of K165 interactions. In bCLC-Ka, K165 interacts closely with the BIM1 sulfonate; in bCLC-Ka_N68D_, K165 is drawn to D68, disrupting BIM1 engagement. (*F*) E72 is positioned to contribute to BIM1 isoform selectivity. The BIM1-bound CLC-Ka structure with binding-pocket residues edited to the CLC-Kb sequence reveals pocket-facing acidic D68 and E72, replacing the neutral N68 and G72 present in CLC-Ka.

Prior studies of benzofuran carboxylates, as well as niflumic and flufenamic acids, established that CLC-K inhibitors typically adopt a non-coplanar arrangement of their two aromatic groups, whereas coplanar geometry is characteristic of activators (8, 20, 25). Based on predicted steric interactions between the chloro substituent and the benzimidazole ring, BIM compounds were similarly expected to adopt a non-coplanar geometry and act as inhibitors (21), and our structures confirm this. Unexpectedly, however, the sulfonate group is not positioned to form a strong interaction with N68 (Fig. 3*A*, right panel), a residue shown by mutagenesis to govern CLC-Ka/Kb isoform selectivity and previously predicted by molecular docking to engage directly with the ligand (21).

### MD simulations illuminate the molecular basis of BIM1 selectivity

Because prior functional data indicate that the identity of residue 68 (N68 in hCLC-Ka, D68 in hCLC-Kb) strongly influences BIM1 potency, our original hypothesis for the selectivity mechanism of BIM1 was that its sulfonate group directly interacts with N68 in hCLC-Ka but is repelled by D68 in hCLC-Kb. However, the distance between N68 and the BIM1 sulfonate in our cryo-EM structure is too far to form a strong interaction. To investigate whether the sulfonate and N68 could form transient interactions not captured in the structure, we ran molecular dynamics (MD) simulations of both wild-type bCLC-Ka and a bCLC-Ka N68D mutant (bCLC-Ka_N68D_) to represent hCLC-Ka and the Ka-to-Kb substitution respectively. Surprisingly, we found no significant difference in the distance between residue 68 and the BIM1 sulfonate between bCLC-Ka and bCLC-Ka_N68D_, and the average distance was about 7-8 Å, too far for any strong interaction to occur (Fig. 3*B*).

While our simulations indicate that no direct BIM1 interaction with N68 is responsible for selectivity, they suggest a different selectivity mechanism. In bCLC-Ka, the BIM1 sulfonate interacts closely with nearby residue K165, whereas in bCLC-Ka_N68D_ the BIM1-K165 distance is increased (Fig. 3 *C,E*). This difference stems from D68 in bCLC-Ka_N68D_ drawing near K165, forming a hydrogen bond that disrupts the ionic K165-BIM1 interaction (Fig. 3 *D,E*). This D68/K165 hydrogen bond occurs much more frequently than does the N68/K165 hydrogen bond in WT bCLC-Ka (67.9% ± 7.0% versus 20.7% ± 5.7% of simulation time, Mann-Whitney U p-value = 0.004). Distance distributions across the full simulation ensemble of bCLC-Ka_N68D_ highlight the larger population of conformations with short K165-D68 interaction distances, in contrast to the smaller population of conformations with short K165-N68 interaction distances in WT bCLC-Ka simulations (Fig. S7). These findings align with a prior CLC-Kb homology model that predicted a D68-K165 interaction (26) and with mutagenesis implicating K165 in BIM1 sensitivity (21). Together, they show that K165 stabilizes BIM1 binding in CLC-Ka far more effectively than in CLC-Kb.

Therefore, the selectivity mechanism for BIM1 is not driven by a direct interaction with residue 68; rather, it arises from altered BIM1 interaction with K165, which can engage BIM1 more readily in hCLC-Ka than in hCLC-Kb. This mechanism also explains why the D68N substitution in hCLC-Kb has only a modest impact on BIM1 potency compared to the reciprocal N68D change in hCLC-Ka (∼3-fold vs ∼13-fold (21)). hCLC-Kb contains an additional negatively charged residue, E72, one helical turn from D68, whereas hCLC-Ka has a neutral glycine at this position. Neutralizing D68 leaves E72 available to sequester K165 away from BIM1 in hCLC-Kb (Fig. *3F*), providing a structural rationale for why the D68N substitution alone does not confer full BIM1 sensitivity. This model predicts that the hCLC-Kb double mutant D68N/E72G should restore BIM1 sensitivity to hCLC-Ka-like levels. Direct functional testing is precluded because neither this mutant nor the more conservative D68N/E72Q variant produced measurable currents in *Xenopus* oocytes. Even so, prior work on negatively charged inhibitors showed that mutating either D68 or E72 enhances potency in hCLC-Kb (27), implicating both residues in modulating inhibitor sensitivity and supporting the overall mechanism of BIM isoform selectivity.

### Conformational flexibility of BIM1 aligns with structure-activity potency trends

In examining the BIM1 binding pose obtained from the cryo-EM structure, we noted that several features of the published BIM SAR data (21) could not be easily reconciled. For instance, introducing bulky phenyl groups to BIM1 at two distinct positions either improved or did not alter BIM1 potency, despite these positions appearing sterically occluded in the modeled pose (Fig. S8 *A,B*). To investigate this discrepancy, we analyzed trajectories of our all-atom MD simulations with BIM1 bound to bCLC-Ka. These simulations revealed that the BIM1 binding pose is dynamic, rather than adopting a single discrete conformation. In particular, the ligand orientation can vary substantially (Fig. S8*B*). Within this ensemble, we identified specific poses that create sufficient space to accommodate bulky substituents, thereby explaining the observed SAR potency trends.

### Extracellular density and atropisomeric poses in the BIM15-bound structure

To further probe the basis of BIM inhibitor selectivity, we determined the structure of bCLC-Ka bound to BIM15 (Fig. S9), which is slightly more potent than BIM1 but lacks CLC-Ka/CLC-Kb selectivity (21). The overall protein conformation and pore profile closely matches the apo and BIM1-bound structures (RMSD=0.5 and 0.4 Å, respectively; Fig. S5*C,E*), but a prominent feature of the BIM15-bound map is the pronounced region of extra density above the ligand toward the extracellular side (Fig. 4*A*). We propose that this density originates from residues in the extracellular I-J loop of bCLC-Ka, which is not sufficiently resolved to model. Focusing on the ligand, we find that BIM15 occupies the binding pocket in two distinct conformations. One conformation matches the pose observed for BIM1. The second maintains the non-coplanar geometry but is rotated such that the chloro substituent projects in the opposite direction (Fig. 4*A*). Following the nomenclature of Bringmann et al. (28), these two poses correspond to the *M*– and *P*-atropisomers of BIM15. Because the two BIM15 poses were so unambiguously resolved, we reexamined the BIM1 density and found that it is best fit by two atropisomeric conformations (Fig. 4*B*), although the difference between its poses is more subtle than for BIM15, whose atropisomers differ by a 17.5° tilt.

**Figure 4.**
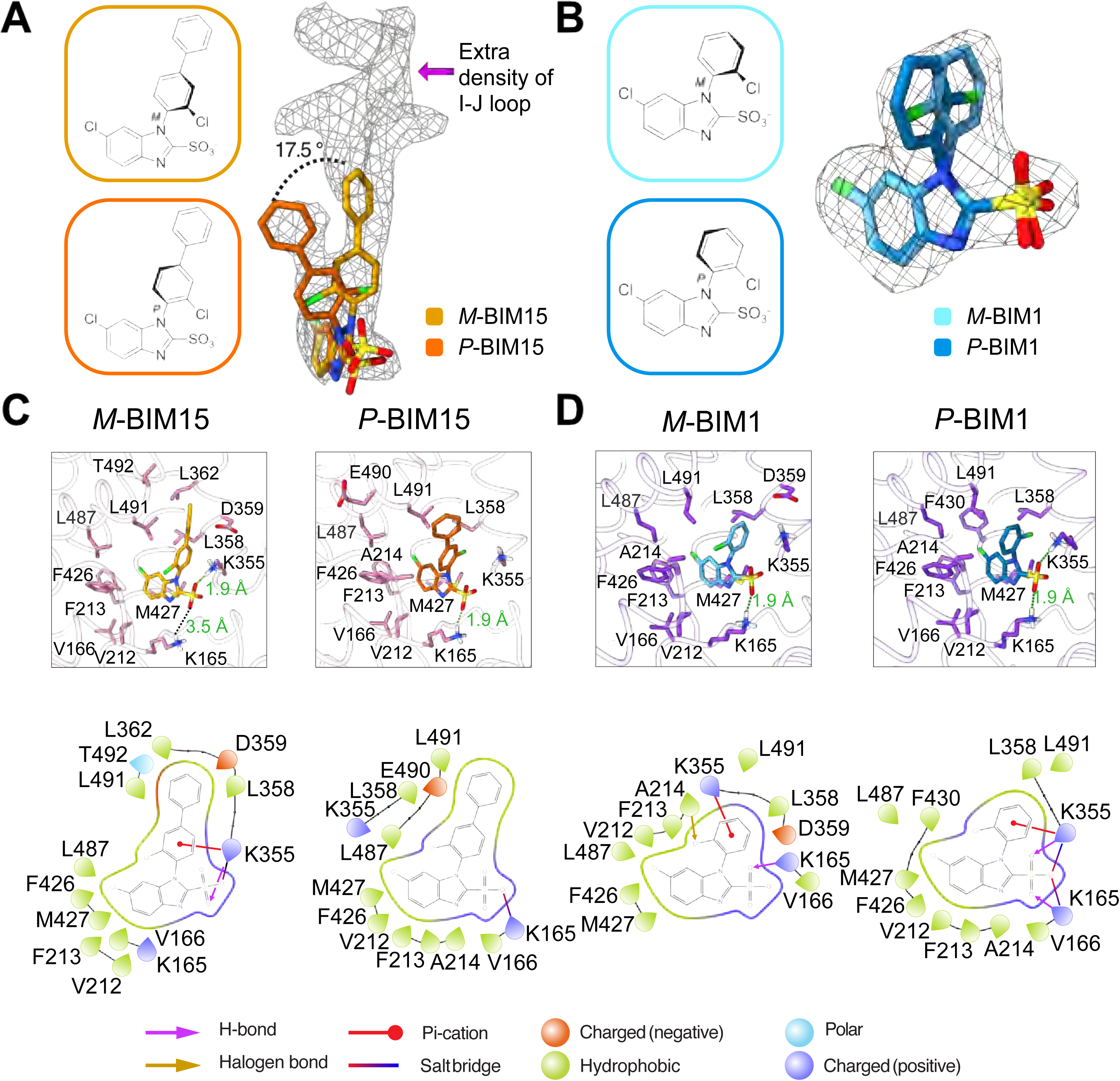
Cryo-EM reveals extra BIM15 density and resolves dual atropisomeric binding modes for BIM15 and BIM1. (*A*) Cryo-EM density for BIM15 overlaid with the molecular models of the two distinct conformations, atropisomers *M*-BIM15 (gold) and *P*-BIM15 (orange). The two atropisomers are 17.5 ° tilted relative to each other. The Q-scores (0.69 for each) indicate that both atropisomers are comparably supported by the cryo-EM density. Extra density appears above the *M*-BIM15 conformation. (*B*) Cryo-EM density for BIM1 overlaid with the molecular models of *M*-BIM1 (cyan) and *P*-BIM1 (blue). The Q-scores (0.72 and 0.74 for *M*– and *P*-BIM1 respectively) indicate that both atropisomers are comparably supported by the cryo-EM density. *M*-BIM1 corresponds to the conformation shown in Fig. 3. (*C*) *Top panels:* Views of the binding site showing Interacting residues within 4 Å of *M*-BIM15 and *P*-BIM15. *Bottom panels:* Corresponding 2D interactions plots generated using the ligand interaction diagram in Maestro (Schrödinger Release 2024.2). (*D*) *Top panels:* Views of the binding site showing Interacting residues within 4 Å of *M*-BIM1 and *P*-BIM1. *Bottom panels:* Corresponding 2D interactions plots generated using the ligand interaction diagram in Maestro (Schrödinger Release 2024.2). The raindrop shape indicates the direction the sidechain is pointing.

### Conserved BIM interactions and the structural basis of BIM15’s lack of isoform selectivity

Structural analysis of the binding site (Fig. 4*C,D*) shows that both BIM1 and BIM15 inhibitors, including their respective atropisomers, are anchored to bCLC-Ka through a conserved polar network and a shared hydrophobic pocket. The sulfonate forms critical hydrogen-bonding interactions with K165 and K355, while the benzimidazole core is stabilized by hydrophobic contacts with V166, V212, F213, A214, F426, M427, and L487, and the chlorobenzene ring interacts with L358 and L491. Although the overall binding modes are conserved between the atropisomers, they show distinct residue-level interactions. For BIM1, the primary distinction is that the *M*-isomer’s chlorobenzene is slightly tilted towards D359 in the upper sub-pocket. Compared to BIM1, BIM15 occupies a larger structural footprint, and its atropisomers differ more markedly, with *M*-BIM15 adopting an upward orientation that enables unique contacts with D359, L362, and T492. Using PDBePISA analysis to quantify the ligand-protein interaction area (see Methods), the BIM1 atropisomers show nearly identical interaction surface areas (325.9 Å² for *M*-BIM1 and 332.4 Å² for *P*-BIM1), whereas BIM15 exhibits a larger atropisomeric difference, with interface surface areas of 391.3 Å² for *M*-BIM15 and 370 Å² for *P*-BIM15. Notably, the *M*-BIM15 interface area is likely underestimated, as the unresolved I-J loop leaves potential loop contacts out of the calculation.

Analysis of the *M*-BIM15 conformation establishes the molecular basis for BIM15’s lack of isoform selectivity between CLC-Ka and CLC-Kb (13). *M*-BIM15 shows a weakened sulfonate-K165 interaction, with an O-H distance of 3.5 Å compared to 1.9 Å in the other BIMs; instead, the sulfonate is stabilized by K355 (1.9 Å O-H distance) (Fig. 4 *C,D*). As a result, the D68 induced displacement of K165 in CLC-Kb has no functional consequence: the absence of a K165 interaction, combined with stabilization by K355, effectively nullifies the K165-dependent selectivity mechanism used by BIM1 and allows BIM15 to inhibit CLC-Kb as efficiently as CLC-Ka. This change in interaction may be driven by BIM15’s engagement with the I-J loop, which sits directly above *M-*BIM15 (Fig. 4A) and appears to pull the inhibitor out of the K165-stabilized position

### Hydrophobicity differences between benzofurans and BIM inhibitors

In developing BIM inhibitors from the less selective but similarly shaped benzofuran series, homology-model docking suggested that CLC-Kb presents a more hydrophobic binding site than CLC-Ka (21). This observation led us to propose that the lower hydrophobicity of the BIM scaffold contributes to its improved selectivity over the benzofurans in the absence of additional stabilizing mechanisms such as engagement of the I-J loop in BIM15 binding. Electrostatic calculations based on the structures determined here support this idea, revealing a similar Ka/Kb difference in the hydrophobicity of the BIM1 binding pocket (Fig. S10). This hydrophobicity-driven contribution to selectivity complements the mechanism outlined above and provides additional context for understanding isoform selectivity of CLC-K channel inhibitors.

### The conformationally flexible I-J loop interacts with bound BIM1 and BIM15

Across all our bCLC-Ka structures – apo, BIM1-bound, and BIM15–bound – we detect density corresponding to the extracellular I-J loop (Fig. 5*A*). While the density in the apo state was sufficient for modeling, albeit with a poor Q-score (Fig. S3*E*), the local resolution from the BIM1-bound, and BIM15–bound maps was too low for confident I-J loop modeling. This region is noteworthy because it appears poised to influence channel gating. In our structures, as well as in the previously reported bCLC-K structure (22), the loop drapes over the pore entryway, positioning it as a plausible gating element. Consistent with this idea, mutagenesis studies have suggested that residues E261 and E278 within the I-J loop form a Ca^2+^-binding site essential for Ca^2+^-dependent gating (29, 30).

**Figure 5.**
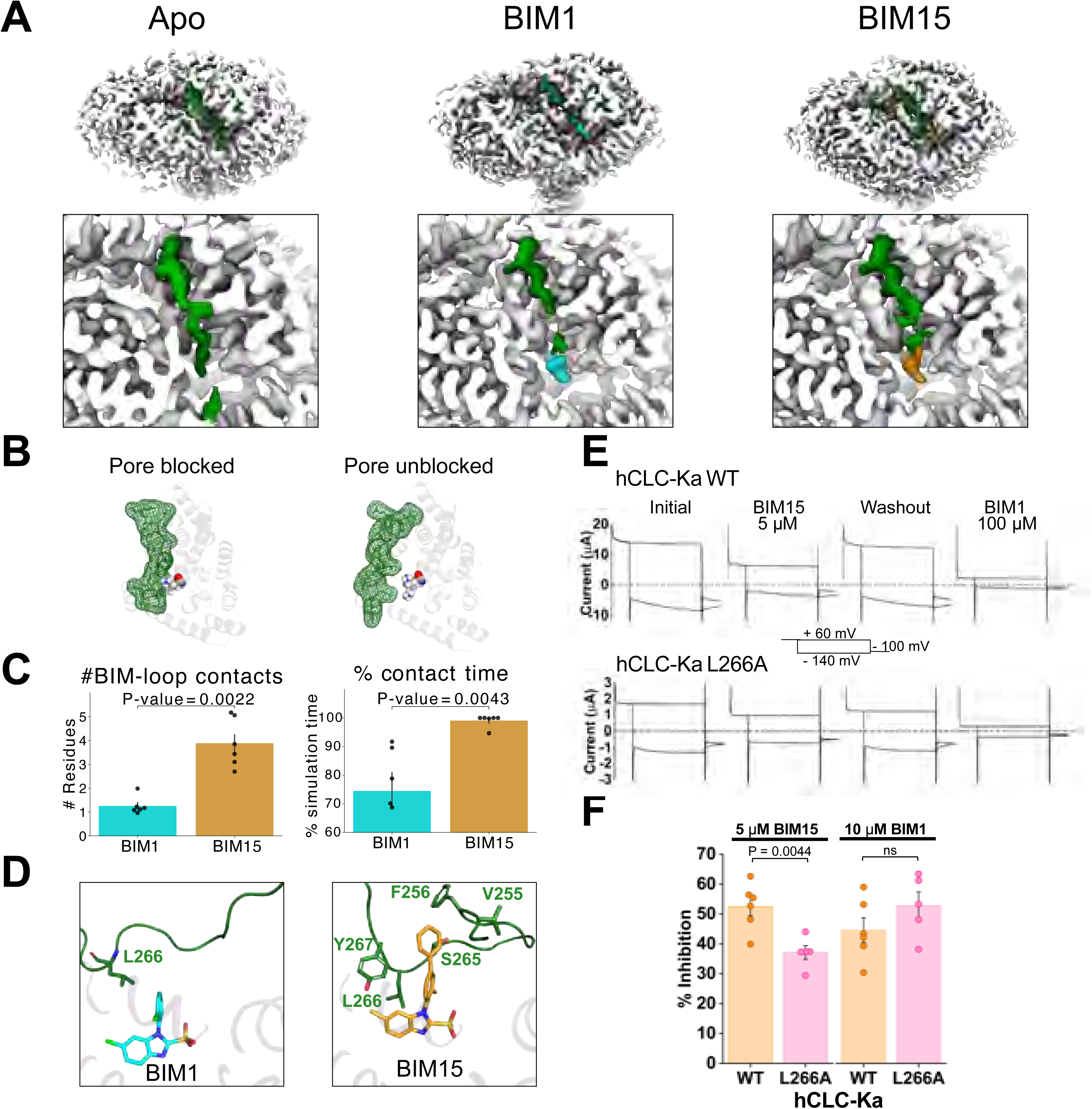
The conformationally flexible I-J loop blocks the pore and interacts with bound BIM1 and BIM15. (*A*) Top views of cryo-EM density for apo, BIM1-bound, and BIM15-bound bCLC-Ka. Upper panels show the full protein; lower panels provide zoomed-in views of the extracellular region. All maps are at the same threshold: 7 standard deviations from the mean. BIM1 and BIM15 densities are shown in cyan and gold respectively. The I–J loop (dark green) spans ∼40 Å between helices and is the only extracellular segment unresolved in our models, supporting its assignment across all three structures. (*B*) Two representative simulation frames from the apo bCLC-Ka simulation, one where the I-J loop blocks the pore, and one where it does not. The I-J loop is displayed as dark green mesh with sticks inside showing the residues. BIM-interacting residues 165, 166, and 355 are displayed as spheres with CPK coloring, marking the position of the channel pore. (*C*) Quantification of contacts to the I-J loop (residues 255-277). *Top:* Average number of loop residues contacting BIM1 or BIM15 (4 Å cutoff) *Bottom:* Contact-time percentage, calculated as the fraction of frames with any loop–ligand contact (4 Å cutoff). For both plots, average values for each individual simulation are shown as black dots, and black bars represent the SEM. P-values were calculated using the non-parametric Mann-Whitney U test. (*D*) MD snapshots showing representative interactions of the I-J loop with BIM1 (top panel) and BIM15 (bottom panel). All residues within 4Å of BIM1/BIM15 are shown. In the representative frames, BIM1 contacts only a single I–J loop residue (L266), whereas BIM15 contacts five; across the full simulation, L266 is engaged during 34% of the BIM1 trajectory versus 96% of the BIM15 trajectory. (*E*) TEVC recordings from oocytes expressing wild-type hCLC-Ka (upper traces) or the I–J loop mutant L266A (lower traces), showing inhibition by 5 µM BIM15. (*F*) Summary of inhibition of hCLC-Ka WT and L266A by 5 µM BIM15 and by 10 µM BIM1. Bars show mean ± SEM; circles represent individual measurements obtained from separate oocytes. Each condition includes data from two independent oocyte batches. P values were determined using unpaired t-tests; for inhibition by 10 µM BIM1, the P value for WT vs L266A is 0.21.

Seeing extra density in the BIM15–bound structure and identifying the unresolved I-J loop as the most plausible source (Figs. 4*A*, 5*A*), we examined MD simulations of bCLC-Ka in which the loop was modeled to the extent permitted by the density. With BIM1 bound, the simulations showed the I-J loop as highly flexible, sampling positions that intermittently blocked access to the BIM1 site (Fig. S11). In apo simulations, the loop was similarly dynamic and moved down into the BIM1 binding pocket, where it intermittently blocked the channel pore (Fig. 5*B*). The loop’s flexibility and its incursions into the BIM site prompted us to ask whether, and how, it interacts directly with bound BIM molecules. To investigate this, we quantified I-J loop contacts with both BIM1 and BIM15. With BIM1, loop residues made only transient and heterogeneous contacts, with no single dominant interaction pattern across the different simulations. In contrast, BIM15 exhibited a recurring interaction with residue L266 in 96% ± 3% of frames and, on average, contacted more loop residues than BIM1 (Fig. 5*C*). These extensive and persistent contacts suggest that the I–J loop contributes to stabilizing BIM15 in a pore-blocking pose; in contrast, the more transient interactions observed with BIM1 do not appear sufficient to provide similar stabilization. The increased contacts likely explain why I-J loop density is continuous with the BIM density in the BIM15-bound cryo-EM map but not in the BIM1-bound structure, and they may contribute to BIM15’s slightly higher potency (55 ± 7% vs. 38 ± 3% inhibition at 5 µM (21)).

Importantly, several features support the reliability of our MD-based conclusion that BIM15 forms more extensive and persistent interaction with the I-J loop than does BIM1. First, this pattern was highly reproducible across six independent simulations for each ligand, with clear separation between BIM1 and BIM15 in both contact frequency and number of loop residues engaged (Fig. 5C). Second, the initial I-J loop coordinates used for simulations were built into the BIM1-bound cryo-EM structure; if imperfections in this modeled starting point biased loop behavior, they would be expected to favor more stable contacts with BIM1, the ligand around which the model was optimized. Instead, we observed the opposite: BIM15 consistently formed more extensive and persistent interactions with the I-J loop. Third, the cryo-EM data themselves align with the simulations, as the BIM15-bound map shows clearer I–J loop density adjacent to the ligand, whereas the BIM1-bound map does not.

To further test the conclusions of our MD simulations and to determine whether the predicted loop engagement has functional consequences, we examined how altering the key interacting residue affects inhibitor sensitivity. We mutated L266, the residue with the highest contact frequency to BIM15 in simulation, and measured BIM1 and BIM15 inhibition. As expected for a model in which extensive loop engagement stabilizes BIM15, BIM15 inhibition was reduced in L266A relative to wild type, whereas BIM1 inhibition was not significantly affected (Fig. 5*E,F*, Fig. S12). The conclusion that I-J loop engagement contributes to potency also aligns with studies showing that I-J loop mutations strongly influence the modulation of CLC-Ka channel activity by niflumic acid, a nonselective chloride-channel inhibitor (31). Representative snapshots of the loop-ligand interactions for BIM1 and BIM15 are shown in Fig. 5*D*.

### The I-J loop mediates Ca²⁺-dependent gating

Our observation that I–J loop movements can block the pore offers a mechanistic connection to earlier work demonstrating that this loop is required for Ca^2+^-dependent gating in CLC-K channels. Extracellular Ca²⁺ activates CLC-K channels in both native and heterologous systems (29, 32–34). In heterologous expression, hCLC-Kb is similarly Ca²⁺ sensitive in *Xenopus* oocytes and HEK293 cells, whereas hCLC-Ka is less Ca²⁺ sensitive in HEK293 cells (23). In native recordings from rat and rabbit, thin ascending limb Cl^‒^ currents carried by CLC-K1 (the paralog of hCLC-Ka) are strongly activated by extracellular Ca²⁺ (32, 34), consistent with the oocyte phenotype and indicating that Ca²⁺-dependent gating observed in oocytes is relevant to the physiological behavior of the renal epithelium. Because Ca^2+^ sensitivity is maximal near physiological concentrations (∼2 mM), and extracellular [Ca²⁺] varies across nephron segments (35, 36), this modulation is well positioned to influence channel activity in vivo.

Foundational work from the Pusch laboratory showed that Ca²⁺ enhances current by influencing channel gating rather than single-channel conductance (29). In these recordings, Ca²⁺ increased open probability without introducing any obvious voltage dependence. Through an extensive mutagenesis campaign, they further demonstrated that neutralizing two conserved acidic residues on the I–J loop abolishes Ca²⁺-dependent activation while leaving permeation intact. Together, these rigorous biophysical experiments identify the I-J loop as the Ca²⁺-responsive element and motivate a structural analysis of how Ca²⁺ binding influences this region.

Given the pronounced flexibility of the I–J loop and its ability to drape over the pore and inhibitor-binding site, we hypothesized that Ca²⁺ binding stabilizes the loop in a conformation positioned away from the pore. To test this idea, we determined the cryo-EM structure of bCLC-

Ka in the presence of 100 mM Ca^2+^ (Fig. S13). The resulting 3.4 Å resolution map shows markedly improved density for the I-J loop (Fig. 6*A*), supported by a higher Q-score (Fig. S13*E*) compared to the 0 mM Ca^2+^ structure (Fig. S3*E*). In the absence of Ca²⁺, the heterogeneous loop density, together with the dynamic movements observed in MD simulations, is consistent with functional studies showing that CLC-Ka retains a non-zero open probability in the absence of Ca² (29). In 100 mM Ca2+, the loop becomes ordered, runs along the subunit interface, and is clearly displaced from the pore (Fig. 6*B*). The pore profile of the Ca²⁺-bound structure closely matches that of the apo channel (Fig. S5*D,E*), and the overall backbone RMSD between the two maps is

**Figure 6.**
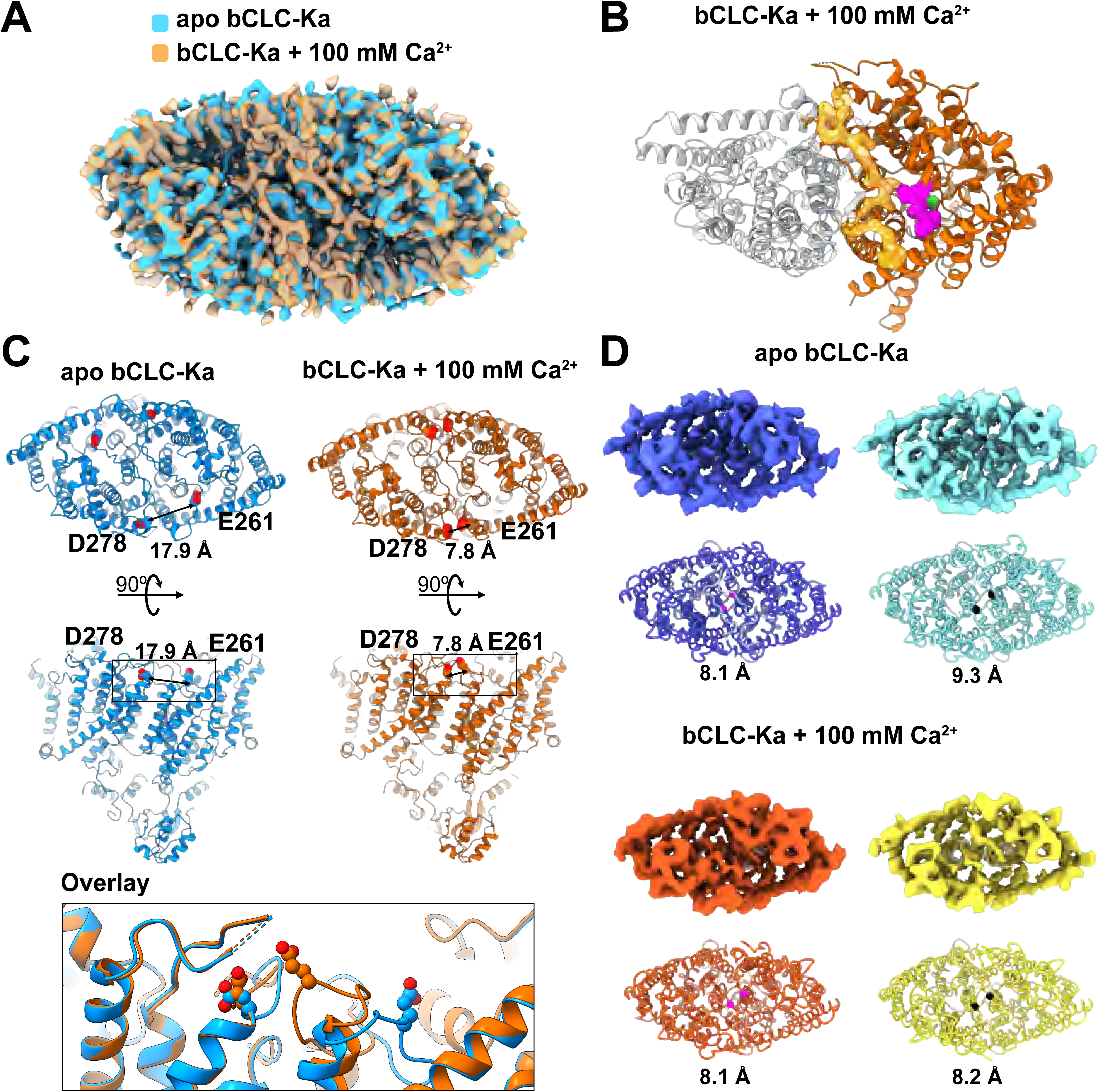
Calcium orders the I-J loop away from the pore. (*A*) Overlay of cryo-EM maps of apo bCLC-Ka (no added Ca²⁺; blue) and bCLC-Ka with 100 mM Ca²⁺ (gold) reveals additional I–J loop density in the Ca²⁺-bound map. Both maps are sharpened and shown at a threshold of 8 standard deviation above means. (*B*) Cryo-EM structure of bCLC-Ka in the presence of 100 mM Ca^2+^. The high-quality density for the I-J loop (orange) supports unambiguous model building of this segment. BIM-interacting residues 165, 166, and 355 are displayed as pink spaced-filled sidechains on the orange subunit, marking the position of the channel pore. (*C*) Ca²⁺ draws E261 and E278 together. In the apo structure (blue), the Cα-Cα distance between the residues is >17 Å, whereas in the 100 mM Ca²⁺ structure (orange), the corresponding Cα-Cα distance contracts to 7.8 Å. (*D*) GMM analysis shows asymmetric subunit motion in apo bCLC-Ka and synchronized, concerted motion in bCLC-Ka with 100 mM Ca²⁺. The top panels show the first (left) and last (right) density maps along the principal motion pathway identified in GMM latent space. The bottom panels show the corresponding structural models, with residue Q494 marked as pink (first snapshot) and black (last snapshot) spheres. In apo bCLC-Ka, the distance between residues changes over the trajectory, whereas in Ca²⁺-bound bCLC-Ka it remains constant, reflecting asymmetric versus synchronous subunit motion.

0.5 Å, indicating that Ca^2+^ binding reorganizes the loop without altering the global architecture. Because 100 mM Ca²⁺ is not fully saturating (29), some residual loop mobility may persist, but any such motion must be slow, as the loop is well resolved in the cryo-EM map. Although Ca²⁺ is not directly visualized in the map, its presence draws the proposed chelating residues, E261 and E278 into proximity, reducing their Cα–Cα distance from 17.9 Å to 7.8 Å (Fig. 6*C*). Nearby residues E259 and E281, previously implicated in tuning CLC-Ka’s calcium sensitivity (30), do not appear positioned to directly coordinate Ca²⁺ with E261 and E278, but may contribute through a general electrostatic effect. Together, these structural changes show that Ca²⁺ stabilizes the I–J loop in a conformation that no longer occludes the pore, providing the structural endpoint of – and independent validation for – the mechanism defined by prior mutagenesis and gating analyses.

### Ca²⁺ enhances coordinated motion in CLC-K channels

These findings raise a natural question: beyond repositioning the loop, does Ca²⁺ binding also reshape the broader conformational dynamics that underlie gating in CLC-K channels? The founding member of the CLC family, CLC-0, exhibits two well-defined gating processes: a fast, independent “protopore” gate in each pore of the homodimer, and a slower “common” gate that opens and closes both pores cooperatively. This dual architecture is thought to apply broadly across the CLC family, although the other homologs have not been characterized with comparable mechanistic resolution. For CLC-K channels, the picture is particularly ambiguous. Because CLC-K channels lack the conserved “glutamate gate” residue that underlies protopore gating in other CLCs, the prevailing assumption has been that the gating observed in single-channel recordings reflects the concerted opening of both pores (1). However, some studies have suggested the presence of an additional protopore-like process (37).

The placement of the Ca²⁺-binding site at the subunit interface suggests that Ca²⁺ binding could promote coupling between the two pores, providing a plausible route for cooperative gating. Our simulations show that the I–J loop occludes only the pore of its own subunit and never crosses over to block the adjacent pore, indicating that any inter-subunit effects must arise from energetic coupling rather than from one subunit directly occluding the other pore. In the Pusch lab’s work, channel activation was described by a four-state allosteric model in which the apparent Ca²⁺ affinity is ∼13-fold higher in the open state than in the closed state, supporting a mechanism in which Ca²⁺ binding at one site energetically favors binding at the other, a hallmark of cooperative gating. Additional evidence for Ca^2+^-dependent cooperativity comes from disease-associated mutations distributed throughout the protein – near the inner and outer pore (V170M, P124L) (38), on an extracellular loop opposite the I–J loop (R351W) (39), and within the cytoplasmic domain (R538P) (40) – all of which modify the channel’s Ca²⁺ dependence. Moreover, transplanting the I–J loop onto CLC-0 alters its common gating in addition to conferring Ca²⁺ sensitivity (30), reinforcing the idea that I-J loop dynamics may be coupled to protein-wide conformational changes.

To probe how Ca²⁺ binding shapes CLC-K channel conformational dynamics, we applied Gaussian Mixture Model (GMM) variability analysis (41) to extract the structural flexibility at the transmembrane domain from our cryo-EM data. While conformational changes were detected in both conditions, the analysis revealed pronounced differences in subunit coupling in bCLC-Ka particles treated with or without Ca²⁺. In the absence of Ca²⁺, the two subunits follow distinct trajectories and move in different directions, consistent with partially independent behavior. Under high Ca²⁺, however, the subunits become dynamically locked and move together in a coordinated fashion (Fig. 6*D*, Fig. S14, supplementary movies). This Ca²⁺-dependent transition from decoupled to concerted motion suggests that Ca²⁺ binding reorganizes the energy landscape and may favor a cooperative gating process, akin to the common gating described in CLC-0 (42, 43) and inferred but not fully defined in other homologs, including CLC-K channels (1, 37). Thus, GMM analysis reveals that Ca²⁺ binding modulates intersubunit coupling, motivating future work to determine whether these Ca²⁺-dependent changes contribute to the poorly understood common gating of CLC-K channels.

### Comparison with previously reported class-2 structure

None of the structures determined in this study match the alternative “class-2” structure reported in the original bCLC-K cryo-EM study (PDB 5TR1). That class-2 structure resembles both the main class-1 structure from the original study (PDB 5TQQ) and the conformations resolved here, but it features a slightly looser dimer interface and a ∼6° tilt between the two transmembrane domains (22). Given similar particle numbers between studies, which reduces the likelihood that sampling depth explains the discrepancy, we hypothesize that the domain-tilted conformation may reflect the presence of detergent and a Fab fragment in the earlier dataset, neither of which was present in our preparations. Thus, the conformations resolved here represent the structural states accessible under our experimental conditions, with a dimer interface similar to that seen in other CLC channels, and they place CLC-Ka’s ligand interactions within a well-defined structural framework.

### Conclusions

CLC-Ka is an attractive therapeutic target for hyponatremia, a common water-balance disorder that remains difficult to manage and often impairs quality of life (44–46). Our structural and computational analyses reveal how benzimidazole inhibitors engage the CLC-Ka binding pocket, showing that interactions with a conserved lysine are central to both inhibition and isoform selectivity, while surrounding residues modulate these contacts. We also identify a dynamic extracellular loop whose movements shape access to the binding pocket and couple to Ca²⁺-dependent gating. Together, these findings unify ligand binding with channel gating and explain how subtle sequence differences translate into distinct pharmacological outcomes, providing a mechanistic framework for designing next-generation CLC-Ka inhibitors and advancing therapeutic strategies targeting renal chloride channels.

## Materials and Methods

### Constructs for CLC expression in *Xenopus* oocytes

Oocyte expression constructs for bovine CLC-K and Barttin were obtained from the MacKinnon laboratory (22). To mimic the human CLC-Ka BIM-binding site, two mutations, S68N and R355K, were introduced into the bovine CLC-K construct. Constructs for human CLC-Ka and Barttin were used as previously described (21). cRNA was generated using linearized DNA templates and the T7 mMessage mMachine Kit (Life Technologies). The human CLC-Kb double mutant D68N/E72G and the human CLC-Ka L266A mutation were generated by GenScript (USA) and verified by whole plasmid sequencing.

### Chemical synthesis

BIM1 and BIM15 were prepared as described in Koster et al. (21)

### Electrophysiological measurements of oocytes expressing CLC-K channels

Electrophysiology experiments for Figure 2 were performed on a fee-for-service basis by NMI Technologie Transfer GmbH. Electrophysiology experiments for Figure 5 were performed in-house. Overall methods and instrumentation were the same for both sets of experiments, with minor differences as noted below.

Oocytes from *Xenopus laevis* (Ecocyte Bio Science, Dortmund, Germany or Xenopus 1 Corp., Dexter, MI) were separated and transferred to a 96-well plate with a conically shaped well bottoms (NUNC) prefilled with Barth’s medium containing 88 mM NaCl, 1 mM KCl, 0.4 mM CaCl_2_, 2.33 mM Ca(NO_3_)_2_, 2.4 mM MgSO_4_, 2.4 mM NaHCO_3_, 5 mM Tris-HCl at pH = 7.6 adjusted with NaOH (NMI experiments) or to a 35-mm culture dish prefilled with Super Barth’s medium (88 mM NaCl, 1 mM KCl, 0.33 mM Ca(NO_3_)_2_, 0.42 mM CaCl_2_. 0.82 mM MgSO_4_, 2.4 mM NaHCO_3_, 10 mM Na-HEPES, 1 mM Na-Pyruvate, 50 mg/ml Gentamycin, pH 7.4) (in-house experiments). Oocytes, were subsequently injected with 5, 10, or 20 ng of hCLC-Ka/Barttin (2:1) RNA mix, or 3.6 ng of RNA of the bCLC-Ka/Barttin (2:1) mix, using a RobooInject System (Multichannel Systems, Reutlingen, Germany; NMI experiments) or manual injection (Drummond Nanoject II; in-house experiments) and stored for 1 day (bCLC-Ka) or 2 – 4 days (hCLC-Ka) after injection.

Automated TEVC recording was performed on a Roboocyte2 setup (Multichannel Systems, Reutlingen, Germany). For experiments shown in Figure 2 (NMI), the oocytes were recorded in ND96 medium (96 mM NaCl, 2 mM KCl, 1 mM MgCl_2_, 1.8 mM CaCl_2_ and 5 mM Na-Hepes at pH = 7.6 adjusted with NaOH). For in-house experiments comparing WT and L266A hCLC-Ka, oocytes were recorded in a high-Ca²⁺ solution (10 mM), which increases currents in both channels and was necessary to obtain reliable signals from the low-expressing L266A mutant. The high-Ca^2+^ solution was prepared following Gradogna et al. (30) and contained 112 mM NaCl, 1 mM MgSO_4_, 10 mM Ca-Gluconate_2_, and 10 mM HEPES, pH = 7.6). The microelectrodes within the measuring heads (Multichannel Systems) were filled with 3 M KCl (NMI experiments) or 1 M KCl, 1.5 M K-Acetate (in-house experiments). Upon oocyte impalement, the holding potential was adjusted to the transmembrane potential (Φ_M_). Oocytes with Φ_M_ > – 10 mV (less negative) were excluded from the measurements, since these values indicate leaky oocytes.

Ion channel currents were assessed in one-minute intervals by a pair of two sequential voltage-clamp recordings that share the same prepulse and tail pulse but differed in their test pulse. Each protocol consistent of a 60-mV prepulse for 50 ms, followed by either a –140-mV test pulse (recording 1) or 60-mV test pulse (recording 2) for 200 ms, and a final tail pulse of –100 mV for 50 ms. For CLC-Ka, currents in response to the –140-mV test pulse served as a quality control because this voltage produced the channel’s characteristic increase in current over the 200-ms test pulse. To quantify this gating behavior, the parameter g was calculated as:

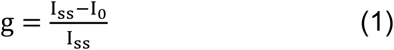

Where I_0_ is the current in the beginning of the –140-mV pulse and I_ss_ the current in the end of the pulse. Recordings with gating parameters below 0.2 were considered to represent non-CLC-Ka currents and were not advanced to inhibitor testing. For bCLC-Ka, which lacks voltage-dependent gating, this metric could not be applied. Instead, oocytes with currents < 800 nA or > 30 µA at 60 mV were not used, as such values indicate either low bCLC-Ka expression or leaky oocytes.

### Calculation of the CLC-K inhibition by BIM1 and BIM15

Inhibition values for BIM1 and BIM15 were determined using a four-step protocol: (1) Initial current (**I**_init_), recorded before inhibitor application. (2) Current recorded after the perfusion of 22 µM BIM1 or 5 µM BIM15 (**I**_c_). (3) Current after the washout of the inhibitors (**I**_WO_) by continuous perfusion of ND96 until I_WO_ returned to 80 – 120 % of I_Init_; experiments that did not meet this washout criterion were not advanced. (4) Current recorded after the perfusion of either 100 µM BIM15 (NMI experiments) or 100 µM BIM1 (in-house experiments) (**I**_L_), which fully inhibit hCLC-Ka currents (Koster et al. 2018). Perfusion was performed at 1 mL/min for 3 min (steps 1, 2 and 4) or > 3 min (step 3). Experiments proceeded only when currents stabilized such that 3 consecutive recordings remained within 5 % of the previous recording.

The inhibition was calculated according to:

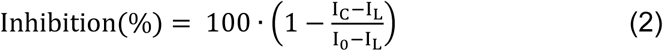

I_0_ is the uninhibited current measured either before inhibitor application (I_init_) or after washout (I_wo_). For each experiment, 2 inhibition values were therefore calculated – one using I_init_ and one using I_WO_. The final inhibition value for that experiment was obtained by averaging these two values. Results are displayed as the mean ± SEM of at least three independent measurements and the statistical significance calculated using an unpaired t-test.

### Protein expression and purification

The bCLC-K construct used for protein production for structural studies was obtained from the MacKinnon laboratory (22). To mimic the human CLC-Ka BIM-binding site, mutations S68N and R355K were introduced into the construct, generating “bCLC-Ka.”. The bCLC-Ka construct was transformed into DH10Bac competent cells (Invitrogen) to generate recombinant bacmid DNA. Purified bacmid was transfected into Sf9 cells using Cellfectin-II (Invitrogen) to produce baculovirus. The baculovirus was further amplified twice in Sf9 cells. The protein was expressed in HEK293S GnTI^−^ cells with the amplified baculovirus. HEK293S GnTI^−^ (ATCC: CRL-3022) cells were cultured in Freestyle 293 medium (Invitrogen) supplemented with 2% fetal bovine serum on a shaker at 37°C in the presence of 8% CO_2_ to a density between 2-3 × 10^6^ cells per mL, then infected with 2% vol/vol baculovirus. After culturing for another 8–16 hours, sodium butyrate was added at final concentration of 10 mM, then further expressed for 48 hours at 37°C before harvest. Cells were harvested by centrifugation and stored at −80°C until use.

2 L of frozen cell pellets were resuspended in resuspension buffer containing 50 mM 4-(2-hydroxyethyl)-1-piperazineethanesulfonic acid (HEPES), pH 7.5, 300 mM NaCl, and protease inhibitor cocktail tablet (MedChem Express), and lysed with 20 strokes of a Dounce homogenizer. Cellular debris were collected by centrifugation at 18,000 rpm for 40 min at 4°C, and then resuspended with resuspension buffer supplemented with 1% n-dodecyl-ß-D-maltoside (DDM) and 0.1% cholesteryl semisuccinate (CHS). After extraction for 2 hours, the lysate was centrifuged at 18,000 rpm for 40 min at 4°C. The clarified lysate was incubated with cobalt resin (TAKARA) for 1 hour at 4°C. Resin was washed with wash buffer containing 50 mM HEPES, pH 7.5, 300 mM NaCl, 10 mM Imidazole, 0.04% DDM, and 0.004% CHS. Purified protein was released from resin with 100 µg HRV 3C protease and incubated at 4°C for overnight. The retrieved protein was concentrated to 0.5 mL with Amicon Ultra (50 kDa cutoff, EMD Millipore) and followed by size-exclusion chromatography (SEC) using a superdex 200 Increased 10/300 chromatography column on an AKTA FPLC system (Cytiva) with buffer containing 20 mM HEPES, 100 mM NaCl, 1 mM DTT, 0.5 mM EDTA, 0.04% DDM, and 0.004% CHS. Protein fractions were pooled, concentrated with Amicon Ultra (50 kDa cutoff, EMD Millipore) to ∼1.5-2.5 mg/mL.

### Cryo-EM sample and grid preparation

All bCLC-Ka cryo-EM structures were determined in Salipro lipid nanoparticles (47) following previously established protocols (48). Purified bCLC-Ka in DDM (1.5–2.5 mg/mL) was mixed with 10 mM POPC, 10 mM POPG, and 1.5 mM Saposin A in incorporation-buffer (50 mM Tris-HCl, pH 7.6, and 150 mM NaCl) to a final volume of 4 mL. The final reaction mixture contained 0.3 mg/mL bCLC-Ka, 1.35 mM POPC, 450 µM POPG, and 120 µM Saposin A. After incubation at 4 °C for 40 min, 50% (w/v) Amberlite XAD-2 (MilliporeSigma) was added for 15 min at 4 °C to remove the detergent. The beads were removed by centrifugation, and the mixture was concentrated to approximately 250 µL using a 10 kDa molecular weight cutoff (MWCO) Amicon centrifugal filter (MilliporeSigma). The sample was further purified by size-exclusion chromatography (SEC) using a Superdex 200 Increase 10/300 GL column (Cytiva) equilibrated in incorporation buffer.

Fractions containing bCLC-Ka in Salipro (eluting at ∼12.3 mL) were pooled and concentrated using a 30 kDa MWCO Amicon filter to a final A280 of 2–4. For ligand-bound states, 1 mM BIM1/BIM15 or 100 mM CaCl_2_ was added and incubated on ice for 2 hours. Prior to grid preparation, 1 mM fluorinated FOS-Choline-8 was added to improve ice thickness and mitigate preferred orientation. For vitrification, 3 µL of the sample was applied to glow-discharged Quantifoil R1.2/1.3 Cu 200-mesh grids, blotted for 3 s with Whatman No. 1 filter paper, and plunge-frozen in liquid ethane using a Vitrobot Mark IV (Thermo Fisher Scientific) at 4 °C and 100% humidity.

### Cryo-EM data acquisition

All cryo-EM data were collected on a Thermo Fisher Scientific Titan Krios cryo-electron microscope operating at 300 kV. The apo bCLC-Ka dataset was acquired using a Falcon 4 direct electron detector without an energy filter. The other three datasets (BIM1, BIM15, and 100 mM Ca^2+^) were collected using a Falcon 4 detector and a Selectris energy filter with a slit width of 10 eV. Data collection parameters are summarized in Table 1. Automated data collection was performed using the EPU software (Thermo Fisher Scientific).

**Table 1.** Cryo-EM data collection, refinement and validation statistics.

|  | bCLC-Ka | bCLC-Ka BIM1 |  | bCLC-Ka BIM15 |  | bCLC-Ka 100 mM Ca<sup>2+</sup> |
| --- | --- | --- | --- | --- | --- | --- |
| **Data collection and processing** |  |  |  |  |  |  |
| Magnification | 96,000 | 165,000 |  | 165,000 |  | 165,000 |
| Voltage (kV) | 300 | 300 |  | 300 |  | 300 |
| Electron exposure (e<sup>-</sup>/Å<sup>2</sup>) | 60 | 60 |  | 60 |  | 60 |
| Defocus range (μm) | -0.8 to -2 | -0.8 to -2 |  | -0.8 to -2 |  | -0.8 to -2 |
| Pixel size (Å) | 0.82 | 0.74 |  | 0.74 |  | 0.74 |
| Symmetry imposed | C2 | C2 |  | C2 |  | C2 |
| Initial particle images (no.) | 3,543,443 | 3,229,270 |  | 421,850 |  | 328,926 |
| Final particle images (no.) | 74,770 | 248,372 |  | 271,313 |  | 191,200 |
| Map resolution (Å) | 3.6 | 2.8 |  | 3.0 |  | 3.4 |
| FSC threshold | 0.143 | 0.143 |  | 0.143 |  | 0.143 |
| Map resolution range (Å) | 2.1 to 11.1 | 1.6 to 8.8 |  | 1.8 to 8.4 |  | 2.2 to 11.3 |
| **Refinement** |  |  |  |  |  |  |
| Initial model used (PDB code) | 5TQQ | P-BIM1 5TQQ | M-BIM1 5TQQ | P-BIM15 5TQQ | M-BIM15 5TQQ | 5TQQ |
| Model resolution (Å) | 3.5 | 2.7 | 2.7 | 2.9 | 2.9 | 3.3 |
| FSC threshold | 0.143 | 0.143 | 0.143 | 0.143 | 0.143 | 0.143 |
| Map sharpening *B* factor (Å<sup>2</sup>) | -116.8 | -110.3 |  | -119.1 |  | -119.5 |
| Model composition |  |  |  |  |  |  |
| Non-hydrogen atoms | 9590 | 9356 | 9356 | 9346 | 9346 | 9592 |
| Protein residues | 1242 | 1208 | 1208 | 1206 | 1206 | 1242 |
| Ligands |  | 6 | 6 | 6 | 6 | 2 |
| *B* factors (Å<sup>2</sup>) |  |  |  |  |  |  |
| Protein | 46.31 | 43.08 | 44.22 | 53.27 | 38.62 | 72.17 |
| Ligand |  | 63.38 | 64.21 | 85.94 | 62.64 |  |
| R.m.s. deviations |  |  |  |  |  |  |
| Bond lengths (Å) | 0.002 | 0.014 | 0.014 | 0.013 | 0.013 | 0.021 |
| Bond angles (°) | 0.494 | 1.892 | 1.867 | 1.880 | 1.773 | 1.663 |
| **Validation** |  |  |  |  |  |  |
| MolProbity score | 1.30 | 1.24 | 1.17 | 0.57 | 1.08 | 1.34 |
| Clashscore | 4.56 | 2.29 | 1.92 | 0.00 | 1.86 | 3.57 |
| Poor rotamers (%) | 1.07 | 0.80 | 0.40 | 0.50 | 0.20 | 1.17 |
| Ramachandran plot |  |  |  |  |  |  |
| Favored (%) | 97.80 | 96.49 | 96.66 | 97.65 | 97.32 | 97.24 |
| Allowed (%) | 2.20 | 3.51 | 3.34 | 2.35 | 2.68 | 2.76 |
| Disallowed (%) | 0.00 | 0.00 | 0.00 | 0.00 | 0.00 | 0.00 |

### Cryo-EM image processing

The complete data processing workflows, including specific details for each sample and dataset, are reported in Figures S3, S4, S7, and S10. For all four datasets (apo, BIM1, BIM15, and 100 mM Ca^2+^), the raw movies were pre-processed (motion correction and CTF estimation) in CryoSPARC Live. Micrographs were curated based on defined criteria, including total motion, CTF fit resolution, and relative ice thickness. Initial particle picking was performed using a blob picker. The resulting particles were pruned through multiple rounds of 2D classification, ab initio reconstruction, and heterogeneous refinement in CryoSPARC. Particles that yielded maps with recognizable protein features were then used for template picking (for the apo dataset) or to train a Topaz particle picker (for the BIM15 and 100 mM Ca^2+^ datasets). Several rounds of heterogeneous refinement (using three classes) were applied to clean the re-picked particle stacks. For the apo dataset, 3D classification without alignment was also performed in RELION, using a mask covering only the protein density for further cleanup. The final reconstructions were generated using non-uniform refinement in CryoSPARC with C2 symmetry imposed.

For the GMM based structure heterogeneity analysis, final particles and their assigned poses from the two Cryo-EM datasets (bCLC-Ka with 0 or 100 mM added Ca^2+^) were imported into EMAN2. Iterative GMM based orientation refinements using a mask focusing on the transmembrane domain (41) were performed to exclude the heterogeneity of the CTD domains and improve the local resolution of the transmembrane domains. The deep learning-based heterogeneity analysis was then applied to the particles with a mask focusing on the transmembrane domains and using information up to 5 Å resolution. The analysis revealed the conformational changes of the region, and the first two eigen trajectories from the analysis were shown for each dataset.

### Model building and structure analysis

A bCLC-K crystal structure (PDB: 5TQQ) was used as an initial atomic model. The initial models were first modified based on our construct (i.e. serine to glutamine and arginine to lysine). These models were rigid-body docked into the cryo-EM map and refined using ISOLDE (49) and PHENIX (50). All atomic models were validated in the PHENIX validation job. Per-residue Q-score (51) was used to assess resolvability of the maps. The I–J loop showed density in all maps, but to varying degrees. In the apo and Ca²⁺-bound structures, the density was sufficiently continuous to support full modeling of the loop, although the low Q-scores in the apo map indicate limited confidence in this region. In contrast, the BIM1– and BIM15-bound maps showed discontinuous density for the I–J loop, and therefore the loop could not be modeled reliably into the deposited structures. For the purposes of MD simulations, we generated loop models as described below, using the available density as a guide where present.

All structure figures were prepared with UCSF ChimeraX (52). 2D interaction plots were generated using the Ligand Interaction Diagram tool in Maestro (Schrödinger Release 2024.2) with a default 4 Å distance cutoff. To prepare the structures, a protein preparation step (including bond order assignment and protonation state assignment at neutral pH) was performed using the Protein Preparation Wizard in Maestro. The ligand-protein interaction areas were calculated with PDBePISA (https://www.ebi.ac.uk/pdbe/pisa/).

Electrostatic surface maps were generated using the Adaptive Poisson-Boltzmann Solver plugin for PyMOL (Schrödinger) and are depicted as surface potentials in Fig. S10. For bCLC-Ka, we calculated potentials directly from the cryoEM structure. For CLC-Kb, we created a “bCLC-Kb” model in PyMOL by mutating all transmembrane domain residues on bCLC-Ka which were identical between bCLC-Ka and hCLC-Ka but different in hCLC-Kb to their hCLC-Kb counterparts. We then calculated potentials on the resulting structure model.

### System setup for molecular dynamics simulations

We performed simulations of bCLC-Ka with five conditions: (1) wild-type apo protein monomer, (2) wild-type monomer with BIM1 bound, (3) wild-type dimer with BIM1 bound, (4) N68D mutant monomer with BIM1 bound, and (5) wild-type BIM15 bound monomer with the I-J loop modeled in. Simulations for (1) through (4) were initiated from a structure of bCLC-Ka bound to BIM1, based on the cryo-EM data reported in this manuscript (specifically, from a model very similar to that presented here but based on an earlier refinement). The I-J loop was modeled into the BIM1-bound structure using the Model Loops function in ChimeraX, which employs the Modeller web server (53). Likewise, simulations for (5) were initiated from an earlier refinement very similar to the structure reported here of bCLC-Ka bound to BIM15. For (1), (2), (4) and (5), one protomer was deleted from the structure using Prime (Schrödinger). For (1), BIM1 was deleted from the pocket using Prime. For (4), N68 was manually changed to an aspartate in Prime. For (5), we aligned the BIM15-bound structure to the BIM1-bound structure, copied over all coordinates for loop residues 256-273, added bonds between the loop and BIM15 residues surrounding the loop, and relaxed the resulting construct using Prime. For each simulation condition, we performed six independent simulations, each between 0.85 and 1 µs in length. For each simulation, initial atom velocities were assigned randomly and independently.

For all simulation conditions, the protein structure was aligned to the Orientations of Proteins in Membranes entry for 5TQQ (bovine CLC-K) using PyMOL. Prime was used to add capping groups to protein chain termini. Protonation states of all titratable residues were assigned at pH 7. Histidine residues were modelled as neutral, with a hydrogen atom bound to either the delta or epsilon nitrogen depending on which tautomeric state optimized the local hydrogen-bonding network. Using Dabble (54), the prepared protein structures were inserted into a pre-equilibrated palmitoyl-oleoyl-phosphatidylcholine (POPC) bilayer, the system was solvated, and sodium and chloride ions were added to neutralize the system at a concentration of 150 mM. The final systems comprised approximately 70,000 atoms, and system dimensions were approximately 80×90×100 Å. Simulations were conducted in the absence of an applied transmembrane voltage.

### Molecular dynamics simulation and analysis protocols

We used the CHARMM36m force field for proteins, the CHARMM36 force field for lipids and ions, and the TIP3P model for waters. Parameters for BIM1 and BIM15 were generated using the CHARMM General Force Field (CGenFF) via the ParamChem web server (55). All simulations were performed using the Compute Unified Device Architecture (CUDA) version of particle-mesh Ewald molecular dynamics (PMEMD) in AMBER20 on graphics processing units (GPUs).

Systems were first minimized using three rounds of minimization, each consisting of 500 cycles of steepest descent followed by 500 cycles of conjugate gradient optimization. 10.0 and 5.0 kcal·mol^−1^·Å^−2^ harmonic restraints were applied to the protein and lipids for the first and second rounds of minimization, respectively. 1 kcal·mol^−1^·Å^−2^ harmonic restraints were applied to the protein for the third round of minimization. Systems were then heated from 0 K to 100 K in the NVT ensemble over 12.5 ps and then from 100 K to 310 K in the NPT ensemble over 125 ps, using 10.0 kcal·mol^−1^·Å^−2^ harmonic restraints applied to protein heavy atoms. Subsequently, systems were equilibrated at 310 K and 1 bar in the NPT ensemble, with harmonic restraints on the protein non-hydrogen atoms tapered off by 1.0 kcal·mol^-1^·Å^-2^ starting at 5.0 kcal·mol^-1^·Å^-2^ in a stepwise fashion every 2 ns for 10 ns, and then by 0.1 kcal·mol^-1^·Å^-2^ every 2 ns for 20 ns. Production simulations were performed without restraints at 310 K and 1 bar in the NPT ensemble using the Langevin thermostat and the Monte Carlo barostat and using a timestep of 4.0 fs with hydrogen mass repartitioning. Bond lengths were constrained using the SHAKE algorithm. Non-bonded interactions were cut off at 9.0 Å, and long-range electrostatic interactions were calculated using the particle-mesh Ewald (PME) method with an Ewald coefficient of approximately 0.31 Å^-1^, and 4th order B-splines. The PME grid size was chosen such that the width of a grid cell was approximately 1 Å. Trajectory frames were saved every 200 ps during the production simulations. The AmberTools17 CPPTRAJ package was used to reimage trajectories (56). Simulations were visualized and analyzed using Visual Molecular Dynamics (VMD) (57) and PyMOL. The GetContacts package was used for calculating the hydrogen bond frequency between residues N/D68 and K165 (https://getcontacts.github.io/). In Fig. S7 to construct the probability distributions for this distance metric, we used trajectory frames from all simulations under each condition and applied a Gaussian kernel density estimator.

## Acknowledgments

We thank Dr. Daniel Collins for synthesizing BIM1 and BIM15. We thank Dr. Alan Pao for sharing his clinical perspective and scientific insight on the physiological context of hyponatremia. We thank Dr. Thomas Jentsch for providing the human CLC-K DNA constructs and Dr. Rod MacKinnon for providing the bovine CLC-K DNA constructs. We thank Dr. Timm Danker (NMI Technologietransfer GmbH) for performing initial Roboocyte2 experiments and for valuable advice on the Roboocyte2 system. We are grateful to Anna Koster, Ria Dinsdale, Catalina Mosquera, and Shwetha Srinivasan for critical reading of the manuscript. This research was supported by the National Institutes of Health grant R01DK128881 (M.M., W.C, and R.O.D.), R01GM150905 (M.C.) and by the Stanford Innovative Medicines Accelerator. The Cryo-EM data was collected at the Stanford-SLAC Cryo-EM Center supported by the National Institute of General Medical Sciences (R24GM154186). A.D. is supported by a postdoctoral fellowship from the American Heart Association, 26POST1564302.

## Author Contributions

C.C., A.S.P., J.K, A.H., R.O.D., W.C, and, M.M designed research; C.C., B.L.S., A.S.P., A.D., J.K., C.N.B., C.F.P., N.T., B.W., C.A.P.W., and M.X. performed research; C.C., B.L.S., A.S.P., J.K., A.D., M.,C., C.F.P., N.T., and C.A.P.W. analyzed data; W.C, R.O.D., and M.M. supervised research; C.C., B.L.S., A.S.P, A.D., J.K., and M.M. wrote the paper; all authors edited the paper.

## Competing Interest Statement

The authors declare no competing interests.

## Classification

Major: Biological Sciences; Minor: Biophysics and Computational Biology

## Figure Legends

**Figure S1.**
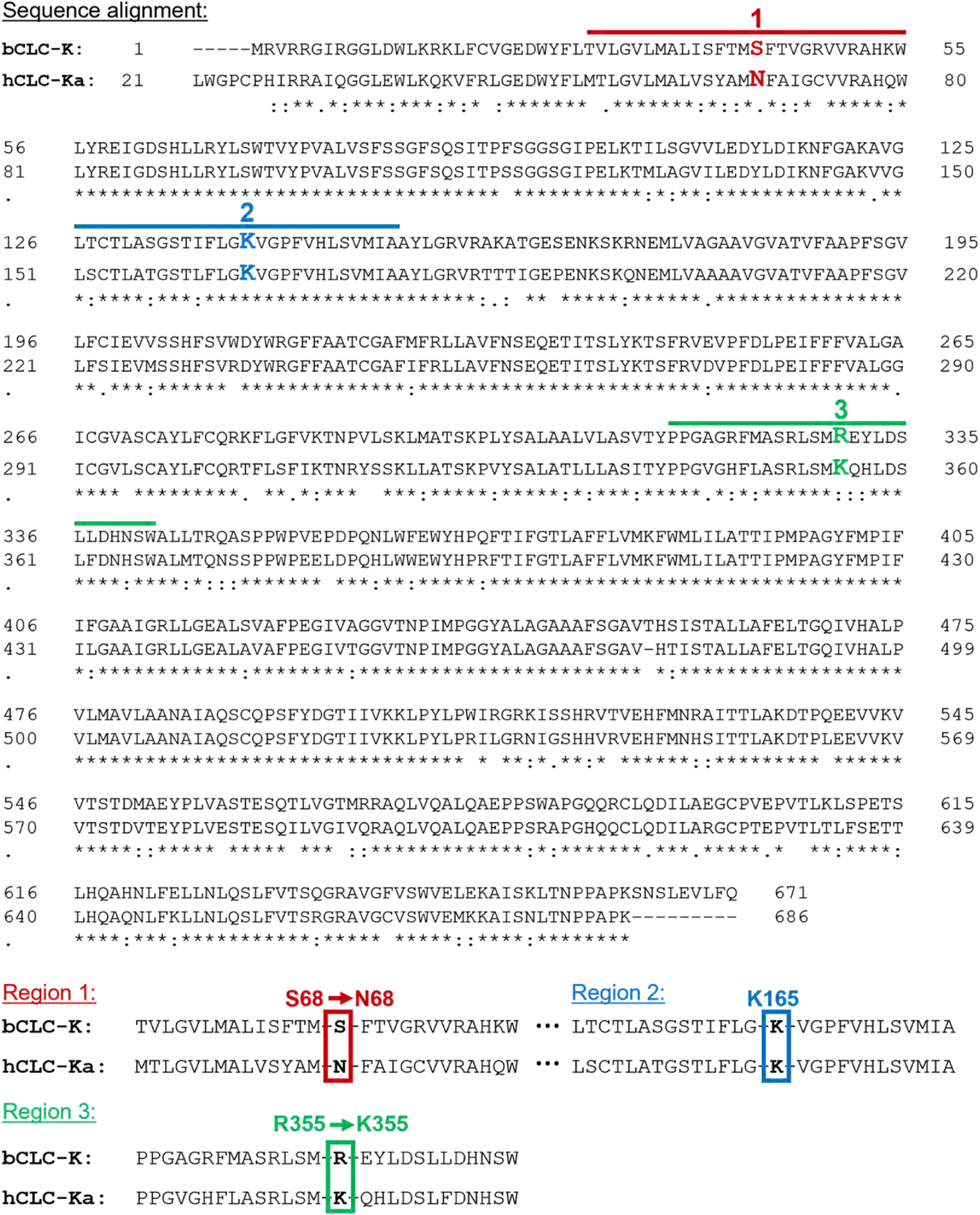
Sequence alignment in the predicted BIM binding pocket.

**Figure S2.**
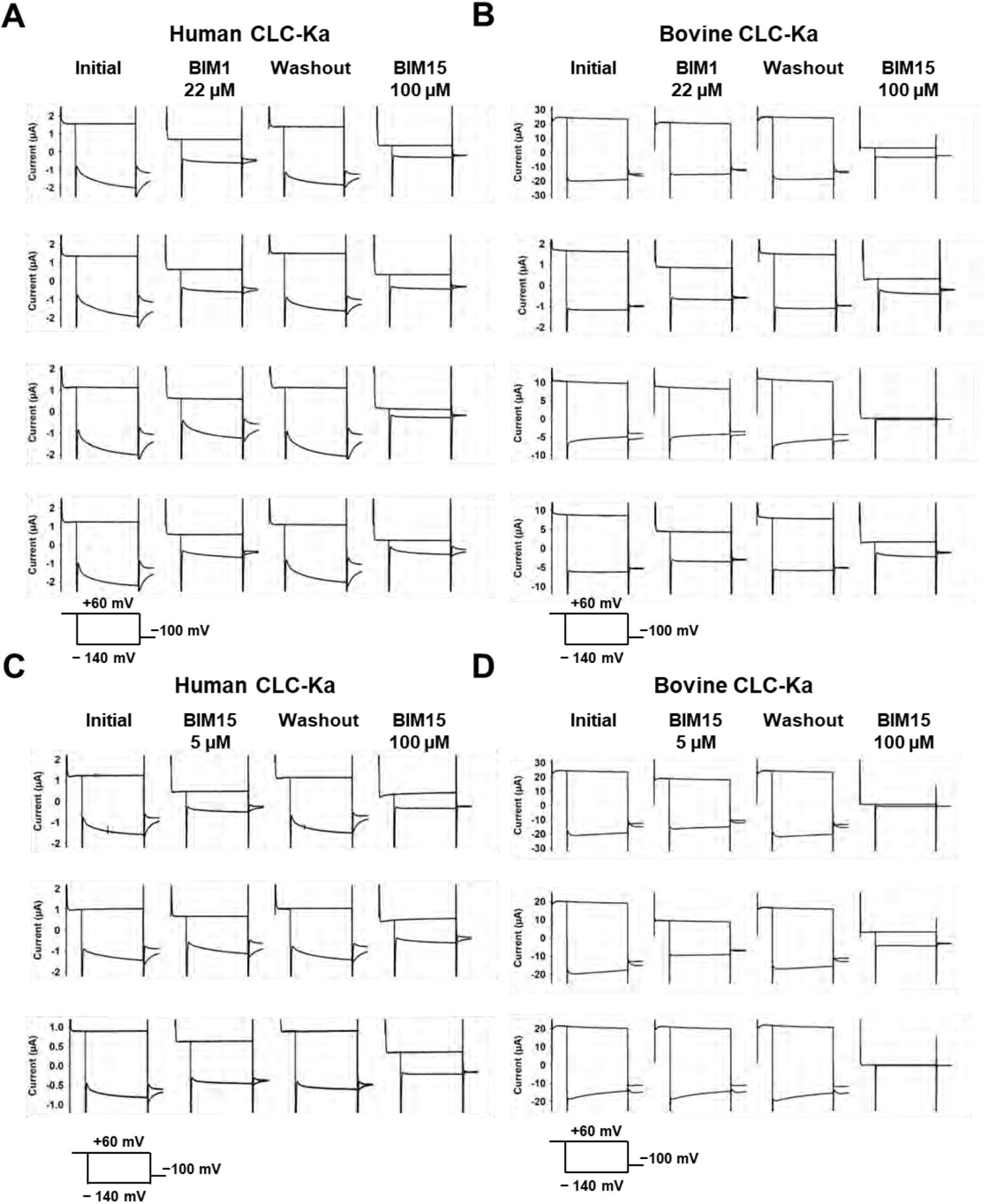
Primary data for TEVC recordings. Data show recordings from oocytes with overexpressed human CLC-Ka. (*A, C*) or bovine CLC-Ka (*B, D*) in the presence of 22 µM BIM1 (*A, B*) or 5 µM BIM15 (*C, D*). Experimental conditions are described in Material and Methods.

**Figure S3.**
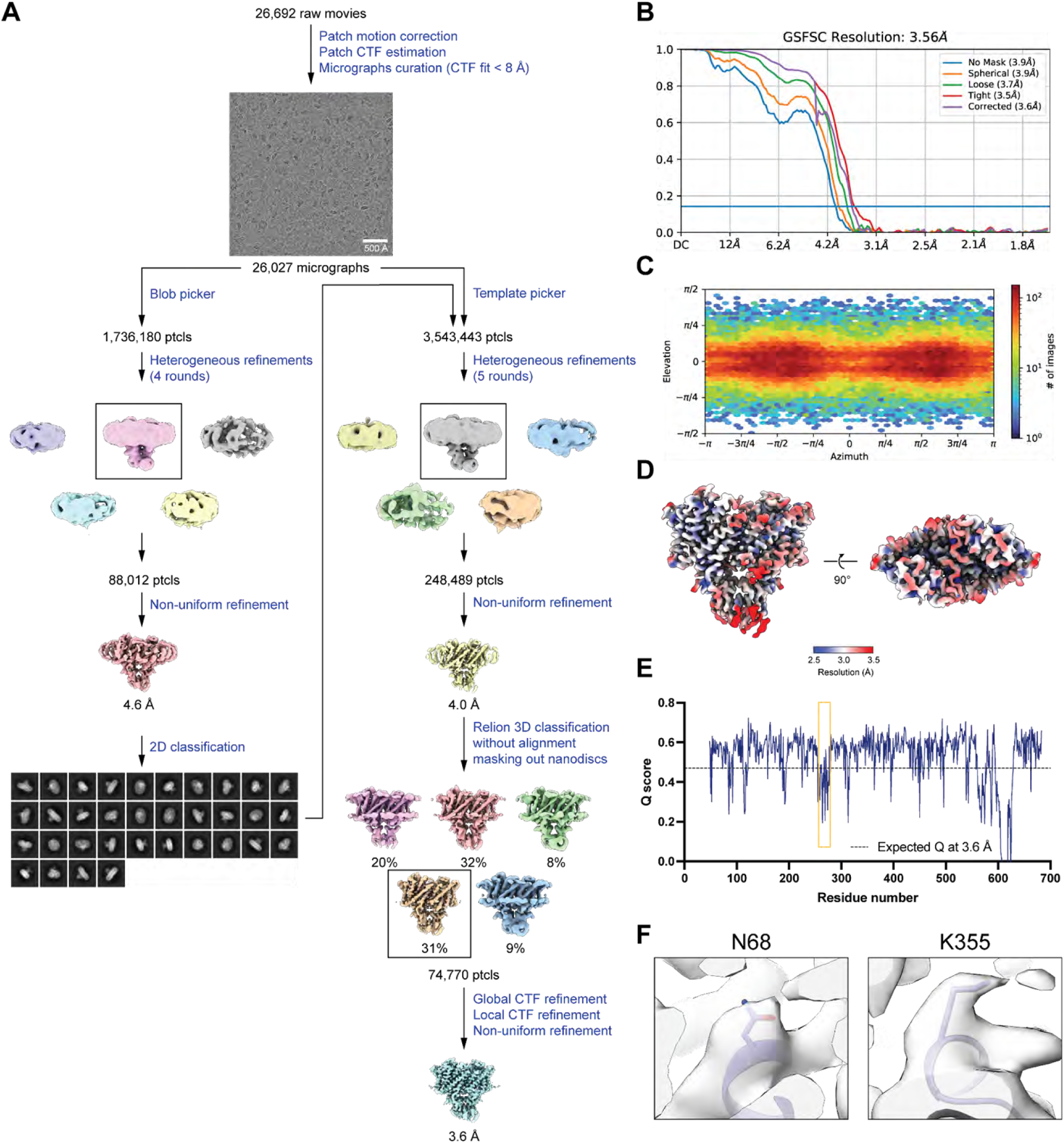
Cryo EM workflow and validation data for bCLC-Ka. (*A*) Cryo-EM data processing workflow. (*B*) Gold-standard FSC curve. The resolution is estimated based on FSC at 0.143. (*C*) Angular distribution plot. (*D*) Local resolution estimation using Locres in cryoSPARC. (*E*) Per-residue Q-score as a function of residue number. The expected Q-score at the map resolution is indicated by dotted line. The I-J loop region is highlighted by orange box. (*F*) The cryo-EM density and molecular model overlay for residue N68 and K355 (the two residues that were mutated to make bovine CLC-K match human CLC-Ka).

**Figure S4.**
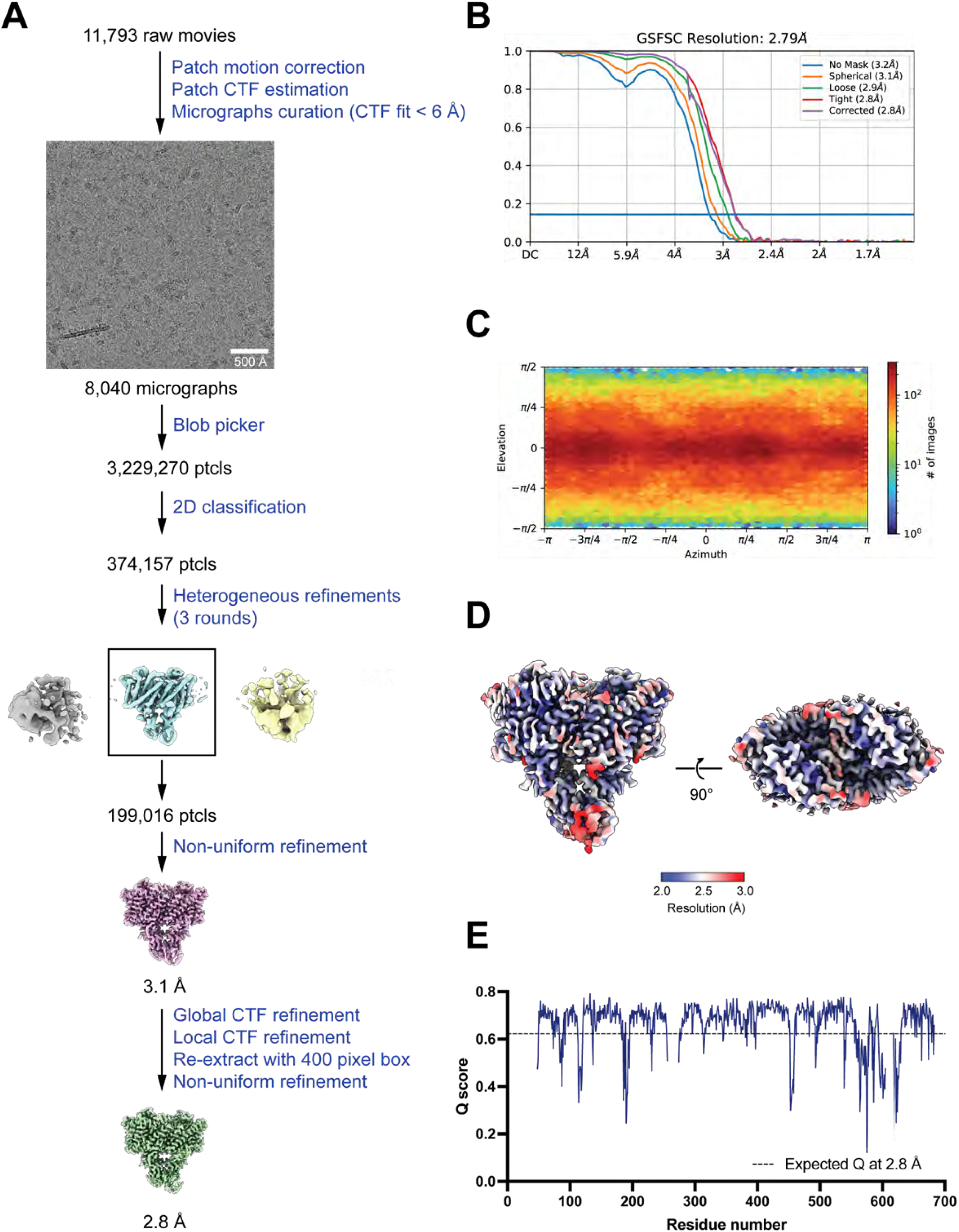
Cryo EM workflow and validation data for bCLC-Ka with BIM1. (*A*) Cryo-EM data processing workflow. (*B*) Gold-standard FSC curve. The resolution is estimated based on FSC at 0.143. (C) Angular distribution plot. (*D*) Local resolution estimation using Locres in cryoSPARC. (*E*) Per-residue Q-score as a function of residue number. The expected Q-score at the map resolution is indicated by dotted line.

**Figure S5.**
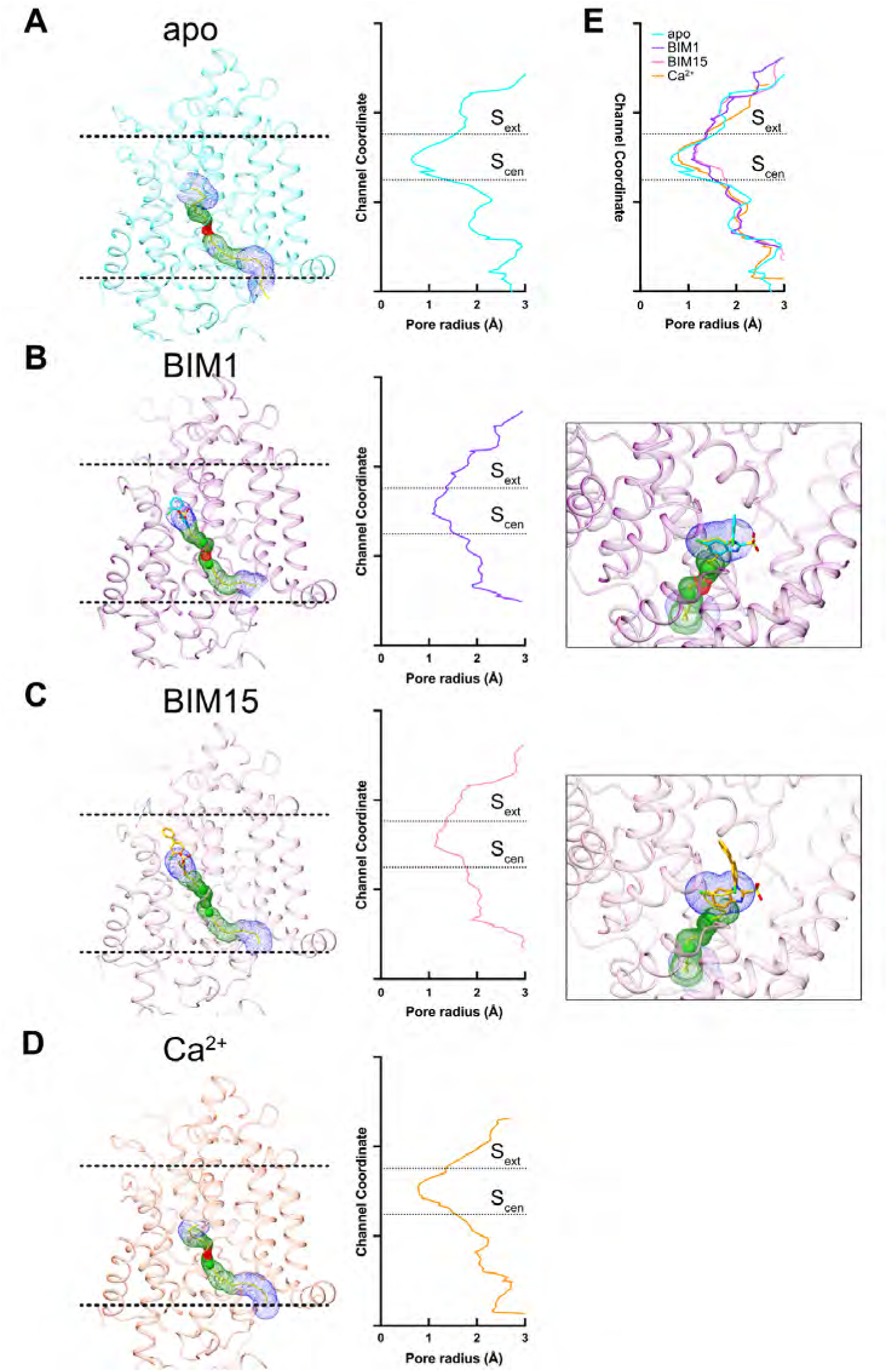
Pore profiles for structures determined in this study. Side views of (A) apo, (B) BIM1-bound, (C) BIM15-bound, and (D) 100 mM Ca2+ bCLC-Ka structures show the pore pathway identified using HOLE; helices B, C, and I are omitted for clarity. The I-J loop in apo structure, the BIM molecules, and the bound chloride ions were removed prior to the HOLE calculation. *M*-BIM1/BIM15 and chloride ions were reintroduced into the final figures to show their positions within the pore pathway. To the right of each structure, pore-radius plots (Å) generated by HOLE illustrate the corresponding profiles, with dashed lines indicating the position of S_ext_ and S_cen_. Close-up views in (B) BIM1– and (C) BIM-15-bound structures highlight the inhibitor molecules occluding the pore. (E) An overlay of all four pore profiles for comparison.

**Figure S6.**
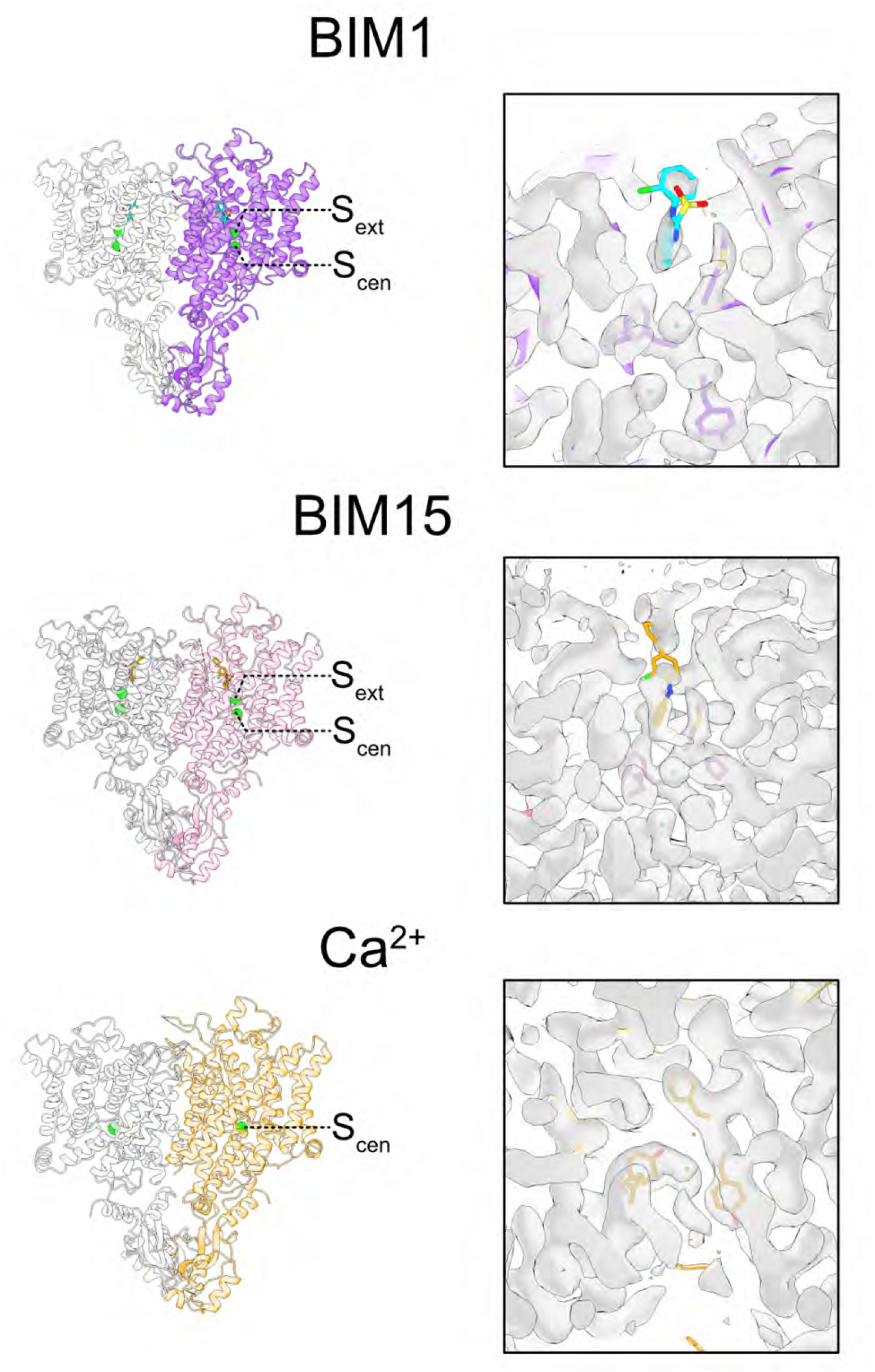
Ion densities for structures determined in this work. The BIM1– and BIM15-bound bCLC-Ka cryo-EM maps display clear densities for Cl^−^ ions at sites S_ext_ and S_cen_ (maps shown at right, models at left), in contrast to the apo maps (not shown), where Cl^−^ densities are not well resolved. The Ca^2+^-bound bCLC-Ka map shows clear density for a Cl^−^ ion at site S_cen_ (map at right, model at left).

**Figure S7.**
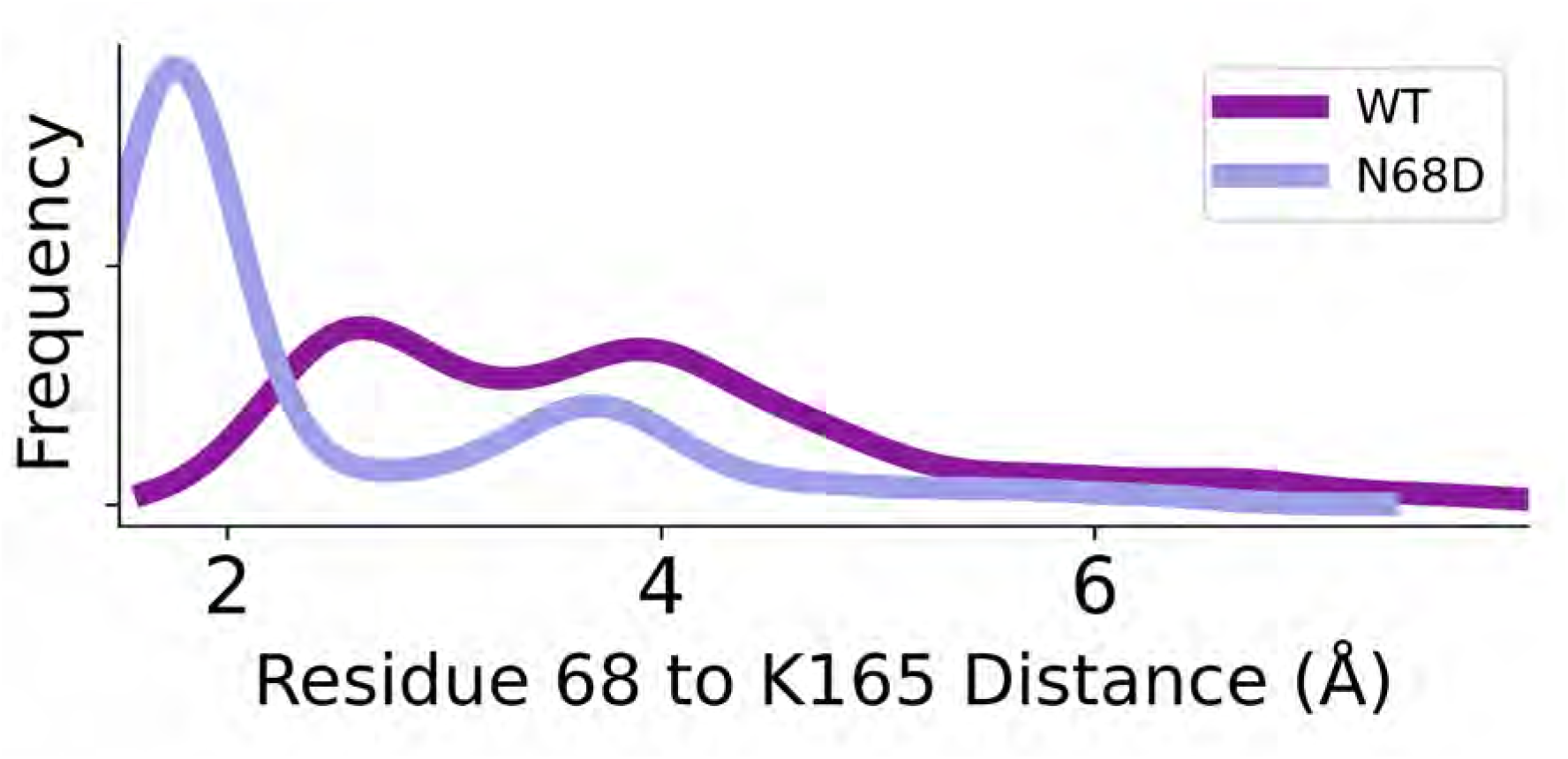
Frequency distribution of residue 68-K165 distance for WT (purple) and N68D (periwinkle) bCLC-Ka in MD simulation. Distributions were calculated using all frames across six independent simulations for each condition (each independent simulation is 0.8-1 μs in length). The distance value was calculated between the closest hydrogen atom on K165 and the closest side chain oxygen/nitrogen of N68 or D68.

**Figure S8.**
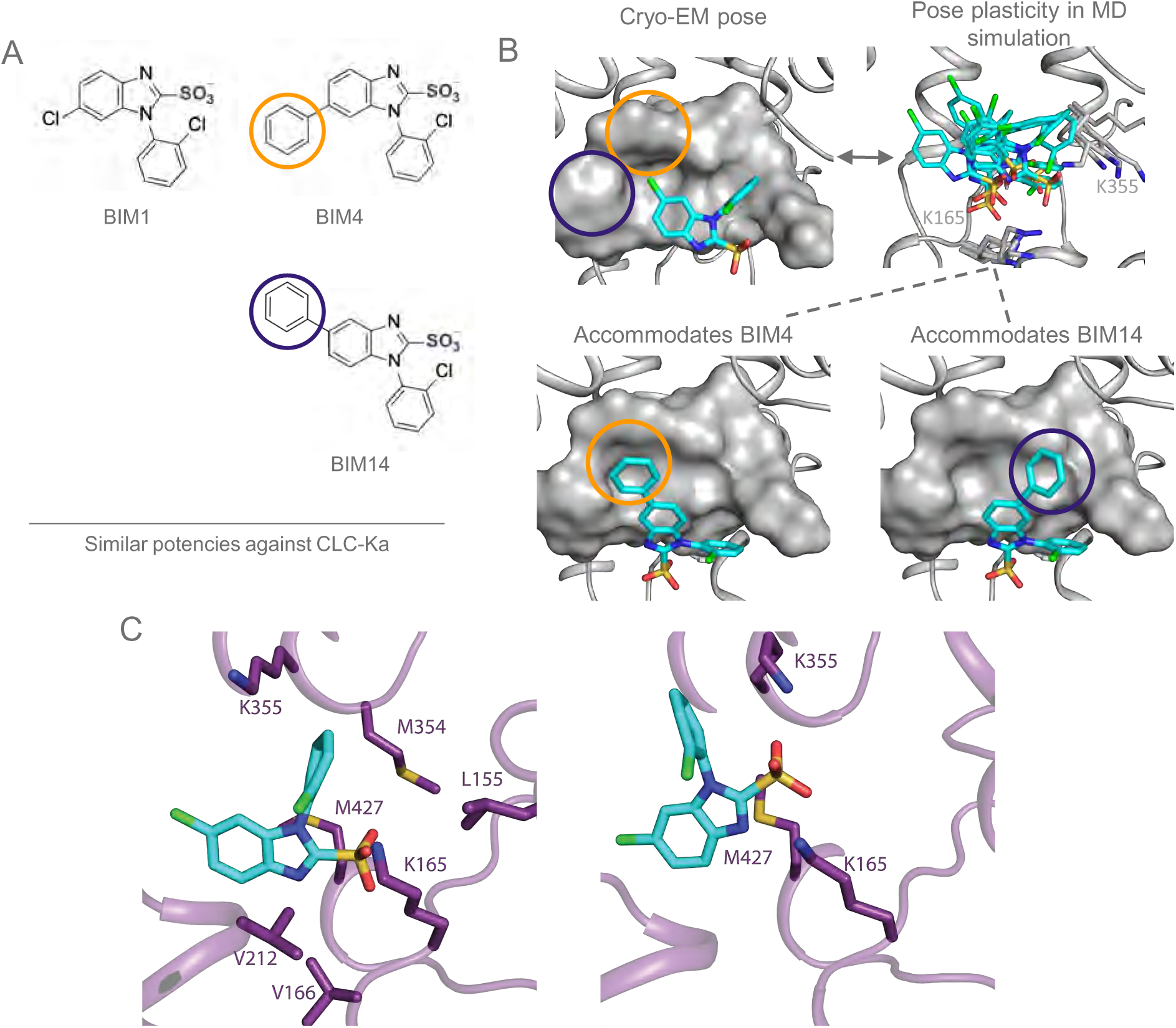
Conformational plasticity of the BIM1 binding pose revealed by molecular dynamics simulations. (*A*) Chemical structures of representative BIM compounds that retain potency but form steric clashes when modeled into the bCLC-Ka structure. BIM4 and BIM14 include additional phenyl groups (highlighted in orange and purple) compared with the parent compound BIM1. (*B*) *Top Left*: Cryo-EM structure of BIM1 bound to bCLC-Ka (gray surface), showing the binding pose (cyan sticks). In this conformation, the additional phenyl groups present in BIM4 and BIM14 would sterically clash with the protein surface (circled). *Top right*: Several frames from MD simulations showing alternative binding poses for BIM1 that could accommodate the additional phenyl groups of BIM4 and BIM14. Residues K165 and K355 are shown as sticks for reference. *Bottom:* Modeled binding poses of BIM4 and BIM14 generated using MD-derived BIM1 poses, illustrating how their phenyl groups could be accommodated in this conformation. (*C*) Representative MD simulation frames showing typical BIM1 interactions with binding pocket residues. All residues within 4Å of BIM1 are shown for each frame. The two poses of BIM1 shown here exhibit different interaction patterns within the binding pocket.

**Figure S9.**
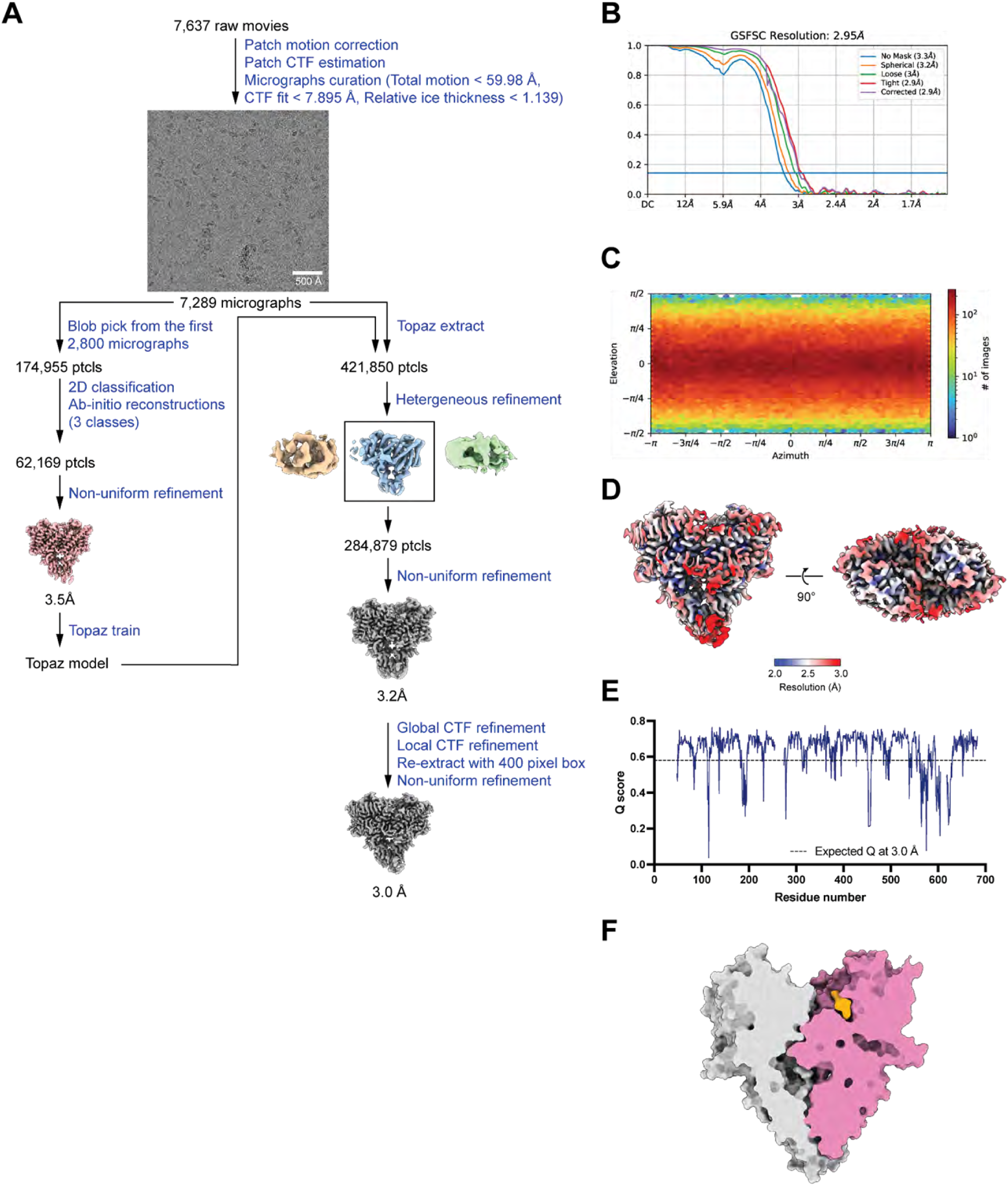
Cryo EM workflow and validation data for bCLC-Ka with BIM15. (*A*) Cryo-EM data processing workflow. (*B*) Gold-standard FSC curve. The resolution is estimated based on FSC at 0.143. (*C*) Angular distribution plot. (*D*) Local resolution estimation using Locres in cryoSPARC. (*E*) Per-residue Q-score as a function of residue number. The expected Q-score at the map resolution is indicated by dotted line. (*F*) Cross-section of the cryo-EM structure of bCLC-Ka with M-BIM15 showing that M-BIM15 directly occludes the chloride pathway.

**Figure S10.**
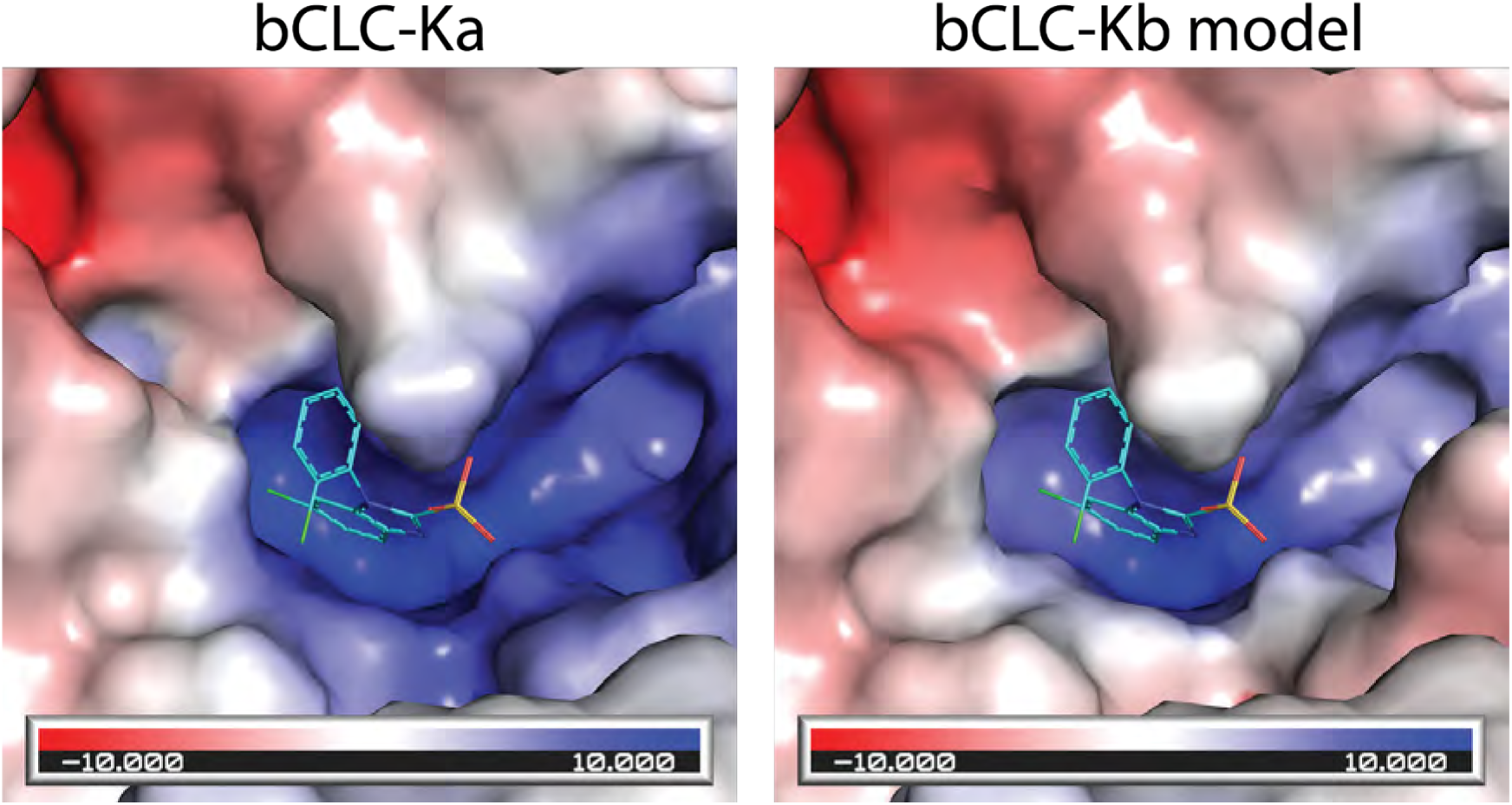
Electrostatic surfaces of bCLC-Ka and a bCLC-Kb model. Electrostatic surface of bCLC-Ka and a bCLC-Kb model, where surface maps are shown on a scale from −10 kT/e (red; negative) to +10 kT/e (blue; positive). BIM1 is shown as cyan lines. These surface maps reveal a more hydrophobic binding pocket in bCLC-Kb versus bCLC-Ka, consistent with previous calculations done on homology models of hCLC-Ka and hCLC-Kb.

**Figure S11.**
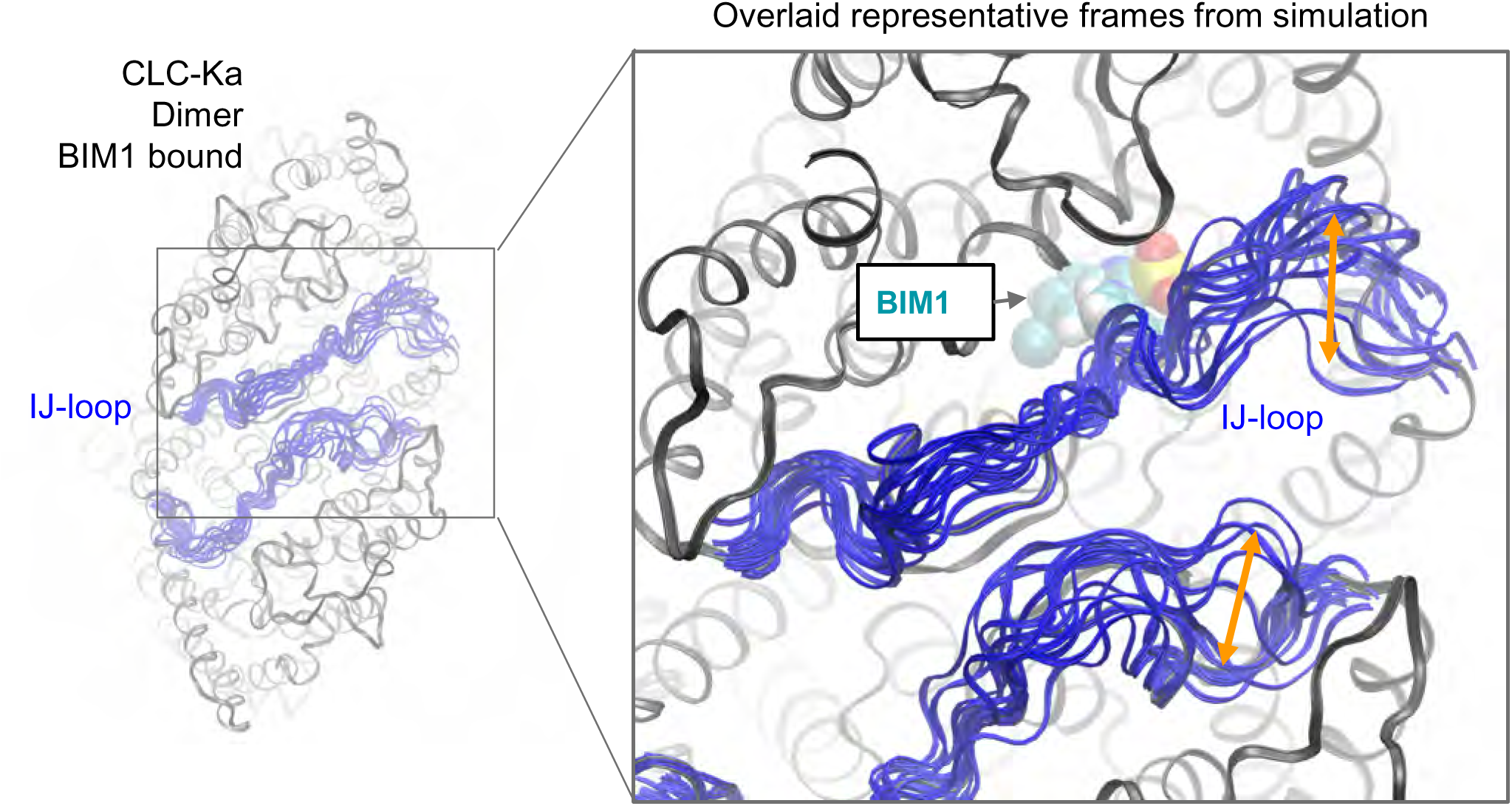
I-J loop flexibility observed in molecular dynamics simulations. Several representative simulation frames of the I-J loop (one frame per every 100 ns) are superimposed onto the experimentally derived structure of bCLC-Ka bound to BIM1. The I-J loop position for each frame is shown as a blue ribbon, while the bCLC-Ka structure is shown as gray ribbons. The experimentally derived pose for BIM1 is displayed for reference and shown as spheres. The I-J loop shows clear flexibility over the course of the simulations, sampling positions that intermittently block access to the BIM1 site.

**Figure S12.**
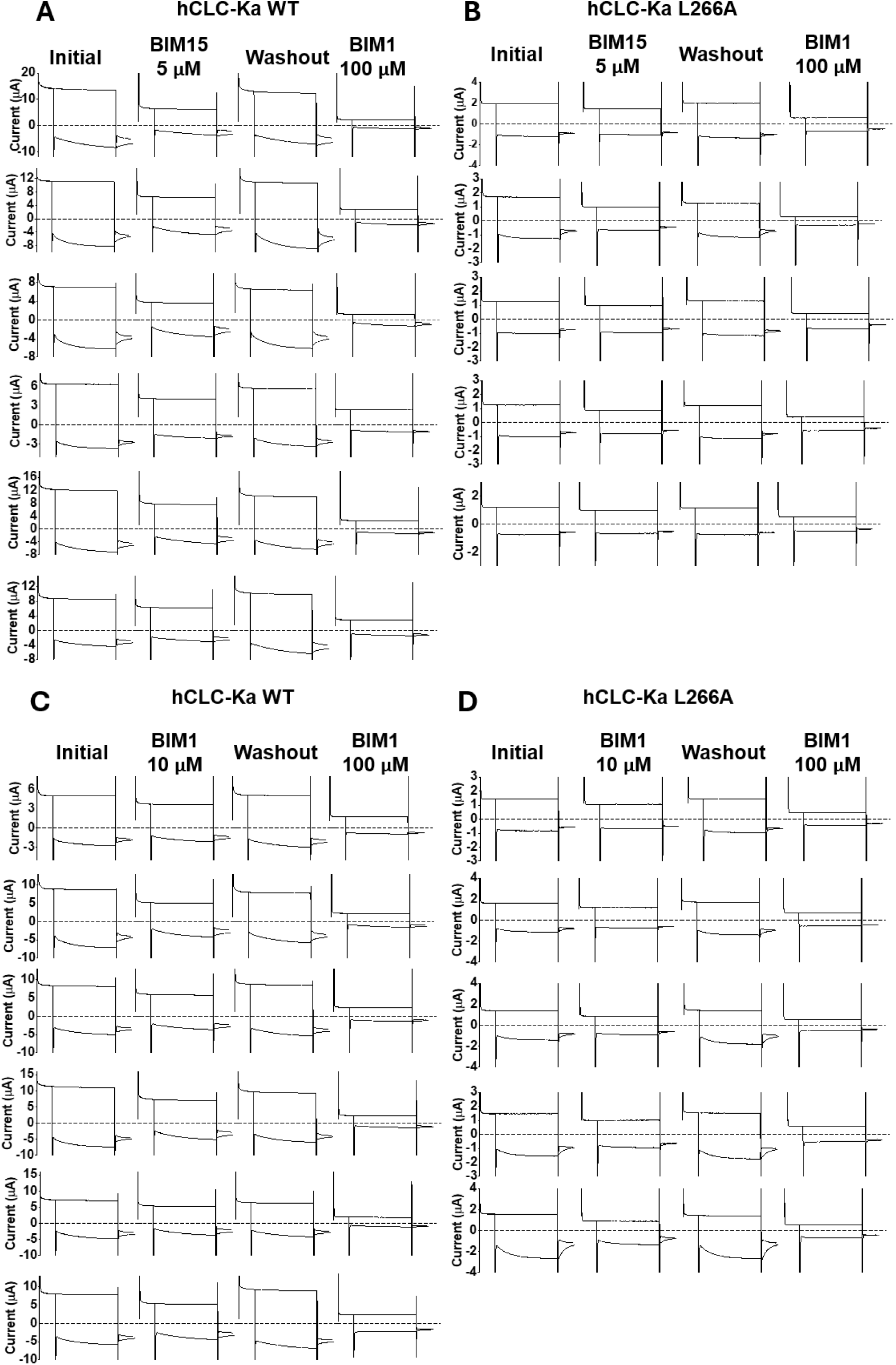
Primary data for TEVC recordings shown in Figure 5. Data show recordings from oocytes with overexpressed WT (A,C) or L266A (B,D) hCLC-Ka channels, with experiments measuring inhibition by 5 µM BIM15 (A,B) or 10 µM BIM1 (C,D). Experimental conditions are described in Material and Methods.

**Figure S13.**
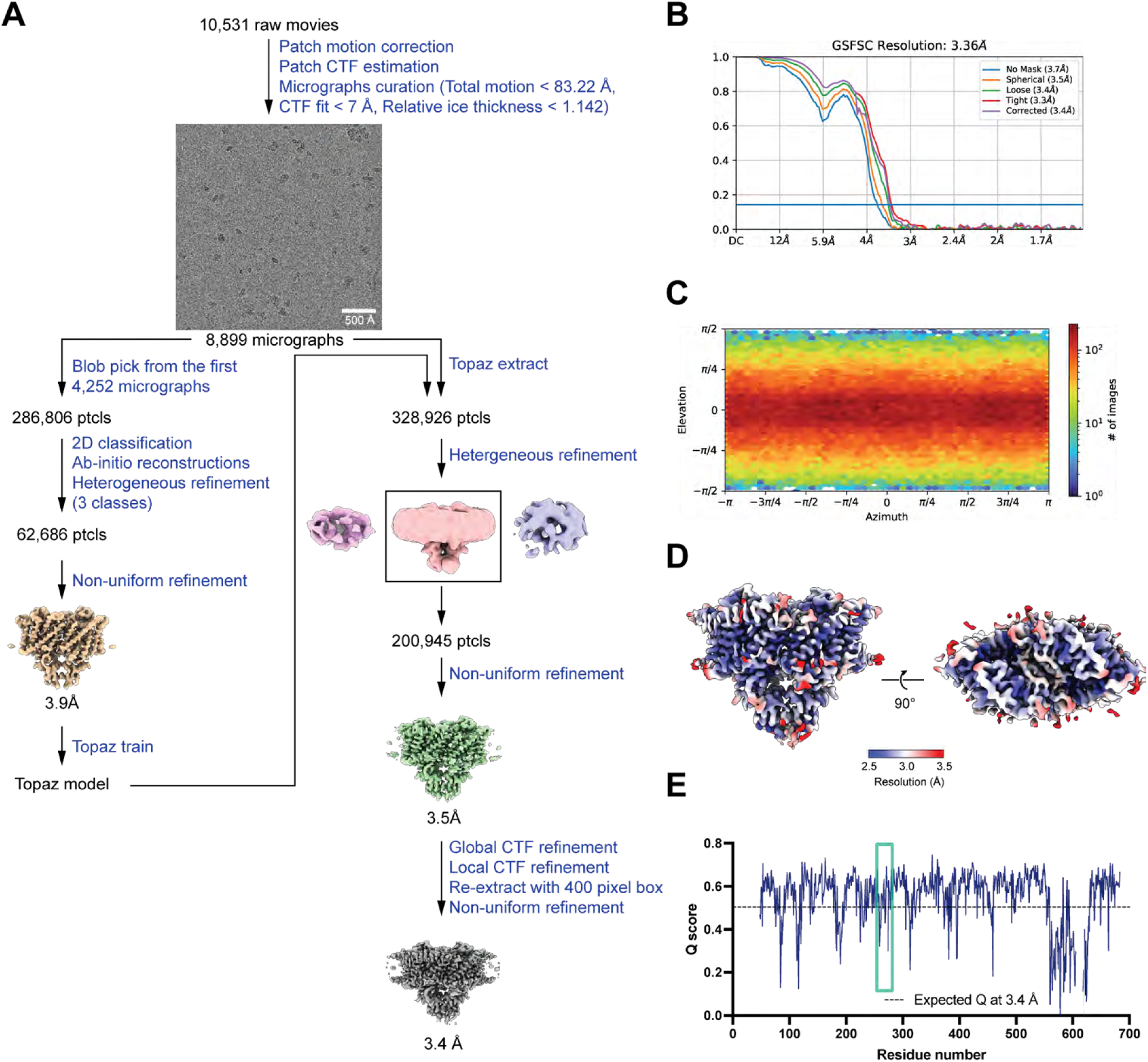
Cryo-EM workflow and validation data for bCLC-Ka with 100 mM Ca^2+^. (*A*) Cryo-EM data processing workflow. (*B*) Gold-standard FSC curve. The resolution is estimated based on FSC at 0.143. (*C*) Angular distribution plot. (*D*) Local resolution estimation using Locres in cryoSPARC. (*E*) Per-residue Q-score as a function of residue number. The expected Q-score at the map resolution is indicated by dotted line. The I-J loop region is indicated by a green box.

**Figure S14.**
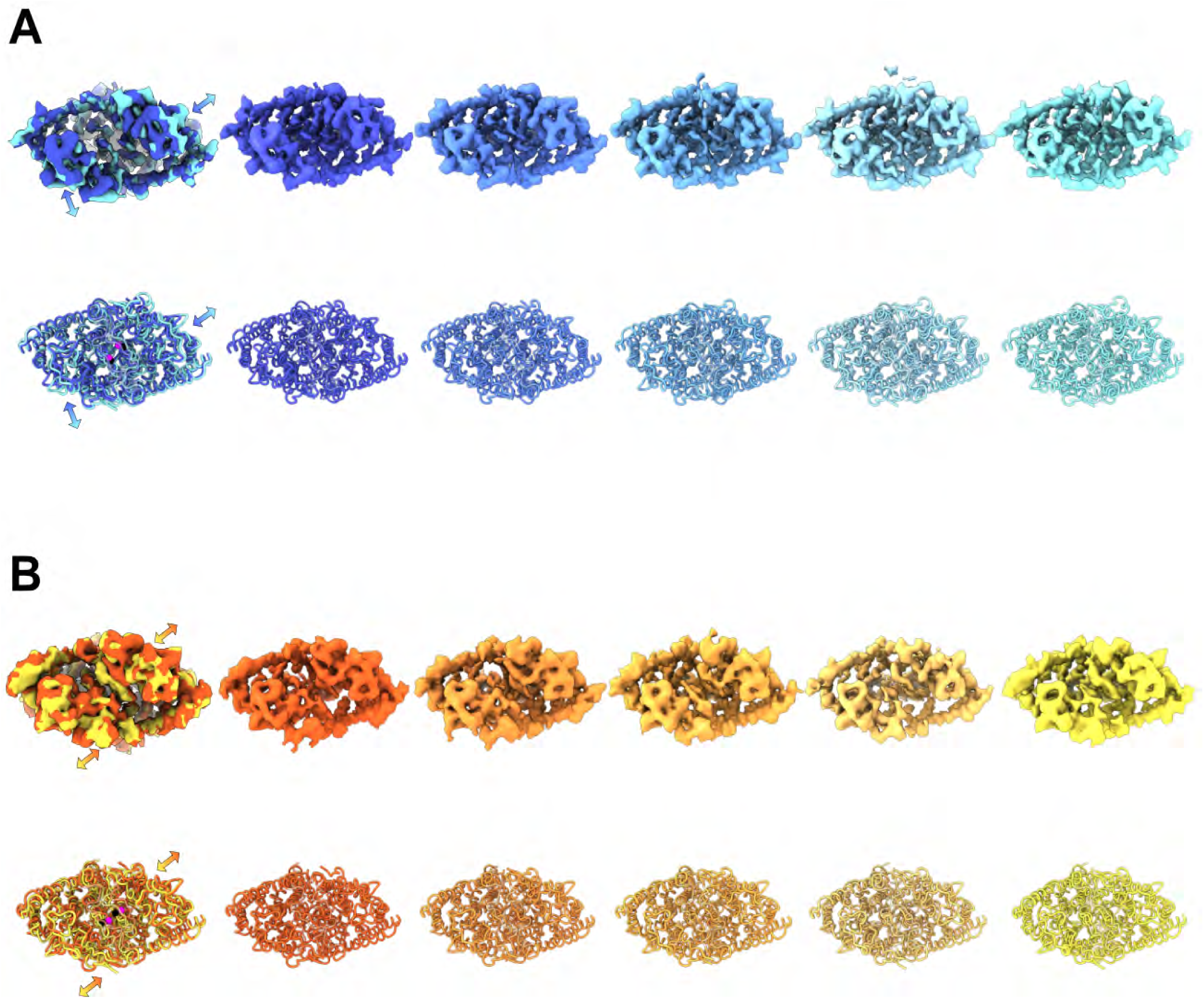
GMM analysis shows asymmetric subunit motion in apo bCLC-Ka and synchronized, concerted motion in bCLC-Ka with 100 mM Ca²⁺. Shown here are the detailed GMM-derived motions that underlie the summary presented in Fig. 6. (*A*) *Apo bCLC-Ka.* The top left panel shows overlays of the first and last density maps along the principal motion pathway identified in GMM latent space. The bottom left panel shows the corresponding overlaid structural models, with arrows indicating the direction of motion. The panels to the right display the five individual maps and models without overlays. (*B*) *Ca²⁺-bound bCLC-Ka.* Panels are arranged as in (*A*).

**Video S1**: [CLC-K_GMM_noCa](https://drive.google.com/file/d/1hhSyCSw9ytPkZqf8nXDWtLbt6xu3HwNg/view?usp=sharing)

**Movie of the asymmetric subunit motion in apo bCLC-Ka revealed by GMM analysis.** The five structural models from the GMM analysis were used to generate the continuous motion using the morph function in ChimeraX. The movie was made in ChimeraX.

**Video S2**: [ClC-K_GMM_Ca](https://drive.google.com/file/d/1rAnBdJjmX8l3vlSXMZtXFH_HwcYMLA4b/view?usp=sharing)

**Movie of the synchronized, concerted motion in apo bCLC-Ka with 100 mM Ca²**⁺ **revealed by GMM analysis.** The five structural models from the GMM analysis were used to generate the continuous motion using the morph function in ChimeraX. The movie was made in ChimeraX.

## Notes

### Competing Interest Statement

The authors have declared no competing interest.

### Summary of Updates

This version has been revised to clarify various points and to add new data (Figure 5E,F; Fig. S12).

## References

1. T. J. Jentsch, M. Pusch, CLC Chloride Channels and Transporters: Structure, Function, Physiology, and Disease. Physiol. Rev. 98, 1493–1590 (2018).

2. A. Picollo, Vesicular CLC chloride/proton exchangers in health and diseases. Front Pharmacol 14, 1295068 (2023).

3. B. K. Kramer, T. Bergler, B. Stoelcker, S. Waldegger, Mechanisms of Disease: the kidney-specific chloride channels ClCKA and ClCKB, the Barttin subunit, and their clinical relevance. Nat. Clin. Pract. Nephrol. 4, 38–46 (2008).

4. C. Fahlke, M. Fischer, Physiology and pathophysiology of ClC-K/barttin channels. Front Physiol 1, 155 (2010).

5. J. Teulon et al., Renal Chloride Channels in Relation to Sodium Chloride Transport. Compr Physiol 9, 301–342 (2018).

6. O. Andrini, D. Eladari, N. Picard, ClC-K Kidney Chloride Channels: From Structure to Pathology. Handb. Exp. Pharmacol. 283, 35–58 (2024).

7. D. B. Simon et al., Mutations in the chloride channel gene, CLCNKB, cause Bartter’s syndrome type III. Nat. Genet. 17, 171–178 (1997).

8. A. Liantonio et al., Kidney CLC-K chloride channels inhibitors: structure-based studies and efficacy in hypertension and associated CLC-K polymorphisms. J. Hypertens. 34, 981–992 (2016).

9. J. P. Kokko, F. C. Rector, Jr., Countercurrent multiplication system without active transport in inner medulla. Kidney Int. 2, 214–223 (1972).

10. J. S. Denton, A. C. Pao, M. Maduke, Novel diuretic targets. Am. J. Physiol. Renal Physiol. 305, F931–942 (2013).

11. M. A. Coppola, M. Pusch, P. Imbrici, A. Liantonio, Small Molecules Targeting Kidney ClC-K Chloride Channels: Applications in Rare Tubulopathies and Common Cardiovascular Diseases. Biomolecules 13 (2023).

12. N. Akizuki, S. Uchida, S. Sasaki, F. Marumo, Impaired solute accumulation in inner medulla of Clcnk1-/– mice kidney. Am. J. Physiol. Renal Physiol. 280, F79–87 (2001).

13. Y. Matsumura et al., Overt nephrogenic diabetes insipidus in mice lacking the CLC-K1 chloride channel. Nat. Genet. 21, 95–98 (1999).

14. R. Birkenhager et al., Mutation of BSND causes Bartter syndrome with sensorineural deafness and kidney failure. Nat. Genet. 29, 310–314 (2001).

15. R. Estevez et al., Barttin is a Cl– channel beta-subunit crucial for renal Cl-reabsorption and inner ear K+ secretion. Nature 414, 558–561 (2001).

16. A. G. Janssen et al., Disease-causing dysfunctions of barttin in Bartter syndrome type IV. J. Am. Soc. Nephrol. 20, 145–153 (2009).

17. R. J. Sepela, J. T. Sack, Taming unruly chloride channel inhibitors with rational design. Proc. Natl. Acad. Sci. U. S. A. 115, 5311–5313 (2018).

18. K. Matulef et al., Discovery of potent CLC chloride channel inhibitors. ACS Chem. Biol. 3, 419–428 (2008).

19. A. Gradogna, M. Pusch, Molecular Pharmacology of Kidney and Inner Ear CLC-K Chloride Channels. Front Pharmacol 1, 130 (2010).

20. A. Liantonio et al., Molecular switch for CLC-K Cl-channel block/activation: optimal pharmacophoric requirements towards high-affinity ligands. Proc. Natl. Acad. Sci. U. S. A. 105, 1369–1373 (2008).

21. A. K. Koster et al., A selective class of inhibitors for the CLC-Ka chloride ion channel. Proc. Natl. Acad. Sci. U. S. A. 115, E4900–E4909 (2018).

22. E. Park, E. B. Campbell, R. MacKinnon, Structure of a CLC chloride ion channel by cryo-electron microscopy. Nature 541, 500–505 (2017).

23. P. Imbrici, A. Liantonio, A. Gradogna, M. Pusch, D. C. Camerino, Targeting kidney CLC-K channels: pharmacological profile in a human cell line versus Xenopus oocytes. Biochim. Biophys. Acta 1838, 2484–2491 (2014).

24. R. Dutzler, E. B. Campbell, R. MacKinnon, Gating the selectivity filter in ClC chloride channels. Science 300, 108–112 (2003).

25. A. Liantonio et al., Activation and inhibition of kidney CLC-K chloride channels by fenamates. Mol. Pharmacol. 69, 165–173 (2006).

26. M. Louet et al., In silico model of the human ClC-Kb chloride channel: pore mapping, biostructural pathology and drug screening. Sci Rep 7, 7249 (2017).

27. A. Picollo et al., Molecular determinants of differential pore blocking of kidney CLC-K chloride channels. EMBO Rep. 5, 584–589 (2004).

28. G. Bringmann et al., Atroposelective synthesis of axially chiral biaryl compounds. Angewandte Chemie 44, 5384–5427 (2005).

29. A. Gradogna, E. Babini, A. Picollo, M. Pusch, A regulatory calcium-binding site at the subunit interface of CLC-K kidney chloride channels. J. Gen. Physiol. 136, 311–323 (2010).

30. A. Gradogna, C. Fenollar-Ferrer, L. R. Forrest, M. Pusch, Dissecting a regulatory calcium-binding site of CLC-K kidney chloride channels. J. Gen. Physiol. 140, 681–696 (2012).

31. A. Gradogna et al., I-J loop involvement in the pharmacological profile of CLC-K channels expressed in Xenopus oocytes. Biochim. Biophys. Acta 1838, 2745–2756 (2014).

32. Y. Kondo, K. Yoshitomi, M. Imai, Effect of Ca2+ on Cl-transport in thin ascending limb of Henle’s loop. Am. J. Physiol. 254, F232–239 (1988).

33. S. Uchida et al., Localization and functional characterization of rat kidney-specific chloride channel, ClC-K1. J. Clin. Invest. 95, 104–113 (1995).

34. R. Sauve, S. Cai, L. Garneau, H. Klein, L. Parent, pH and external Ca(2+) regulation of a small conductance Cl(-) channel in kidney distal tubule. Biochim. Biophys. Acta 1509, 73–85 (2000).

35. M. M. Mupanomunda, B. Tian, N. Ishioka, R. D. Bukoski, Renal interstitial Ca(2+). Am. J. Physiol. Renal Physiol. 278, F644–649 (2000).

36. C. Prot-Bertoye, L. Lievre, P. Houillier, The importance of kidney calcium handling in the homeostasis of extracellular fluid calcium. Pflugers Arch. 474, 885–900 (2022).

37. M. Fischer, A. G. Janssen, C. Fahlke, Barttin activates ClC-K channel function by modulating gating. J. Am. Soc. Nephrol. 21, 1281–1289 (2010).

38. O. Andrini et al., CLCNKB mutations causing mild Bartter syndrome profoundly alter the pH and Ca2+ dependence of ClC-Kb channels. Pflugers Arch. 466, 1713–1723 (2014).

39. Y. Yu et al., Identification and functional analysis of novel mutations of the CLCNKB gene in Chinese patients with classic Bartter syndrome. Clin. Genet. 77, 155–162 (2010).

40. G. Q. Martinez, M. Maduke, A cytoplasmic domain mutation in ClC-Kb affects long-distance communication across the membrane. PLoS ONE 3, e2746 (2008).

41. M. Chen, S. J. Ludtke, Deep learning-based mixed-dimensional Gaussian mixture model for characterizing variability in cryo-EM. Nat. Methods 18, 930–936 (2021).

42. T. J. Jentsch, Discovery of CLC transport proteins: cloning, structure, function and pathophysiology. J. Physiol. 593, 4091–4109 (2015).

43. C. Miller, In the beginning: a personal reminiscence on the origin and legacy of ClC-0, the ‘Torpedo Cl(-) channel’. J. Physiol. 593, 4085–4090 (2015).

44. H. Sumi, N. Tominaga, Y. Fujita, J. G. Verbalis, C. G. and the Electrolyte Winter Seminar, Pathophysiology, symptoms, outcomes, and evaluation of hyponatremia: comprehension and best clinical practice. Clin Exp Nephrol 29, 134–148 (2025).

45. I. Portales-Castillo, R. H. Sterns, Allostasis and the Clinical Manifestations of Mild to Moderate Chronic Hyponatremia: No Good Adaptation Goes Unpunished. Am. J. Kidney Dis. 73, 391–399 (2019).

46. P. T. Brinkkoetter et al., Impact of Resolution of Hyponatremia on Neurocognitive and Motor Performance in Geriatric Patients. Sci Rep 9, 12526 (2019).

47. J. A. Lyons, A. Boggild, P. Nissen, J. Frauenfeld, Saposin-Lipoprotein Scaffolds for Structure Determination of Membrane Transporters. Methods Enzymol. 594, 85–99 (2017).

48. C. H. Chien et al., An Adaptable Phospholipid Membrane Mimetic System for Solution NMR Studies of Membrane Proteins. J. Am. Chem. Soc. 139, 14829–14832 (2017).

49. T. I. Croll, ISOLDE: a physically realistic environment for model building into low-resolution electron-density maps. Acta Crystallogr D Struct Biol 74, 519–530 (2018).

50. P. V. Afonine et al., Real-space refinement in PHENIX for cryo-EM and crystallography. Acta Crystallogr D Struct Biol 74, 531–544 (2018).

51. G. Pintilie et al., Measurement of atom resolvability in cryo-EM maps with Q-scores. Nat. Methods 17, 328–334 (2020).

52. E. F. Pettersen et al., UCSF ChimeraX: Structure visualization for researchers, educators, and developers. Protein Sci. 30, 70–82 (2021).

53. N. Eswar et al., Comparative protein structure modeling using Modeller. Curr Protoc Bioinformatics Chapter 5, Unit-5 6 (2006).

54. R. M. Betz, v2.6.3. 10.5281/zenodo.836914 (2017).

55. K. Vanommeslaeghe, Shen, N., Polani, N., Fan, Y., Ghosh, J., Herath, C., Marru, S., Pierce, M., Pamidighantam, S, Sheetz, M., Mackerell, (2012) ParamChem force field parametrization engine: Initial guess generation and dihedral parameter optimization. in American Chemical Society.

56. D. R. Roe, T. E. Cheatham, 3rd, PTRAJ and CPPTRAJ: Software for Processing and Analysis of Molecular Dynamics Trajectory Data. Journal of chemical theory and computation 9, 3084–3095 (2013).

57. W. Humphrey, A. Dalke, K. Schulten, VMD: visual molecular dynamics. J. Mol. Graph. 14, 33–38, 27-38 (1996).

